# Multi-scale tissue fluorescence mapping with fibre optic ultraviolet excitation and generative modelling

**DOI:** 10.1101/2022.12.28.521919

**Authors:** Joel Lang Yi Ang, Ko Hui Tan, Alexander Si Kai Yong, Chiyo Wan Xuan Tan, Jessica Sze Jia Kng, Cyrus Jia Jun Tan, Rachael Hui Kie Soh, Julian Yi Hong Tan, Kaicheng Liang

## Abstract

Cellular imaging of thick samples requires physical sectioning or laser scanning microscopy, generally incompatible with high-throughput requirements. We developed fibre optic microscopy with ultraviolet (UV) surface excitation (FUSE), a portable, quantitative fluorescence imaging platform for thick tissue that substantially advances prior UV excitation approaches with illumination engineering and computational methods. Optic fibres delivered <300nm light with directional control, enabling unprecedented 50X widefield imaging on thick tissue with sub-nuclear clarity, and 3D topography of surface microstructure. Generative modelling of high-magnification images using our normalising flow architecture FUSE-Flow (open-source) enhanced low-magnification imaging by variational inference. Comprehensive validation comprised multi-scale fluorescence histology compared with standard H&E, and quantitative analyses of senescence, antibiotic toxicity, and nuclear DNA content in tissue models via efficient sampling of entire murine organs by thick slices up to 0.4×8×12mm and 1.3 million cells per surface. This technology addresses long-standing laboratory gaps for high-throughput studies for rapid cellular insights.

**Teaser:** Large-field functional cellular insights into thick tissue with generative AI enables accelerated decision-making

## 1 Introduction

Comprehensive spatial insights from tissue and organs at the ‘mesoscopic’ centimetre scale are of growing interest. Fluorescence imaging of large intact samples is a key enabler in protein expression studies [1] and organ-level spatial transcriptomics [2]. Confocal [3], multiphoton microscopy [4], and light-sheet microscopy with tissue clearing [5] are important techniques today but are costly and complex commercial offerings, usually available as shared infrastructure in privileged scientific organisations. An uncomplicated and economical microscope for rapid cellular imaging of thick, large samples could potentially scale to every lab bench, biotechnology outfit, and less-privileged hospital. This concept has been a long-standing challenge in surgical pathology [6] and is emerging in life science with stringent requirements on quantitative imaging, molecular specificity, and compatibility with laboratory assays [7]. Specifically for tissue, a fast analogue to conventional hematoxylin and eosin (H&E) histology is needed for thick or intact samples [8]. Label-free microscopy of biological samples based on optical coherence tomography [9] or quantitative phase imaging [10] are attractive for their simplicity in sample preparation, but the lack of molecular multi-colour contrast hampers image interpretation and completely precludes multiplexed readouts.

The use of deep ultraviolet (DUV) illumination for fluorescence excitation is a recent promising approach for thick-sample microscopy without lasers or scanning. DUV light penetrates unsectioned tissue superficially, providing optical sectioning, and can simultaneously excite multiple fluorophores using a single wavelength [11, 12]. Also known as microscopy with ultraviolet surface excitation (MUSE), this technique’s simplicity has been previously highlighted for its potential in intra-operative imaging [13, 14], although demonstrations in life science have been limited [15, 16]. Previous MUSE designs typically delivered DUV illumination directly from light-emitting diodes (LEDs) at an oblique angle. These designs required specialised long-working-distance objective lenses, adding to cost and size. Demonstrated magnifications in the literature were limited up to 10X, likely due to insufficient optical axial sectioning achieved by direct illumination.

We propose fibre optic microscopy with UV surface excitation (FUSE), featuring a new fibre optic illumination strategy (Fig. 1a) that enables magnifications up to an unprecedented 50X, 0.55 NA (Fig. 1c), surpassing capabilities of prior MUSE designs. Leveraging precise angular control of the fibre optic illuminators, near-horizontal (up to *∼*87° from vertical) illumination minimised the effective penetration depth and enhanced optical sectioning for high magnification (Fig. 1b). The fibre optic delivery also enabled oblique illumination for low-cost objective lenses with conventionally small working distances (down to *∼*1 *mm*), more efficient heat management of light sources at a distance, and notably, micro-scale 3D topography based on sequential omnidirectional illumination and photometric stereo reconstruction [17]. Our 30×30 *cm* platform is integrated with stepper motors, precision stages, and a colour camera for capturing large fields-of-view (Fig. 1d). Illumination was synchronised strictly to camera capture for minimisation of DUV exposure to tissue samples. Virtual H&E recolouring [18, 19] can be performed to suit standard histopathology appearance for ease of clinical interpretation (Fig. 1e).

**Fig. 1:**
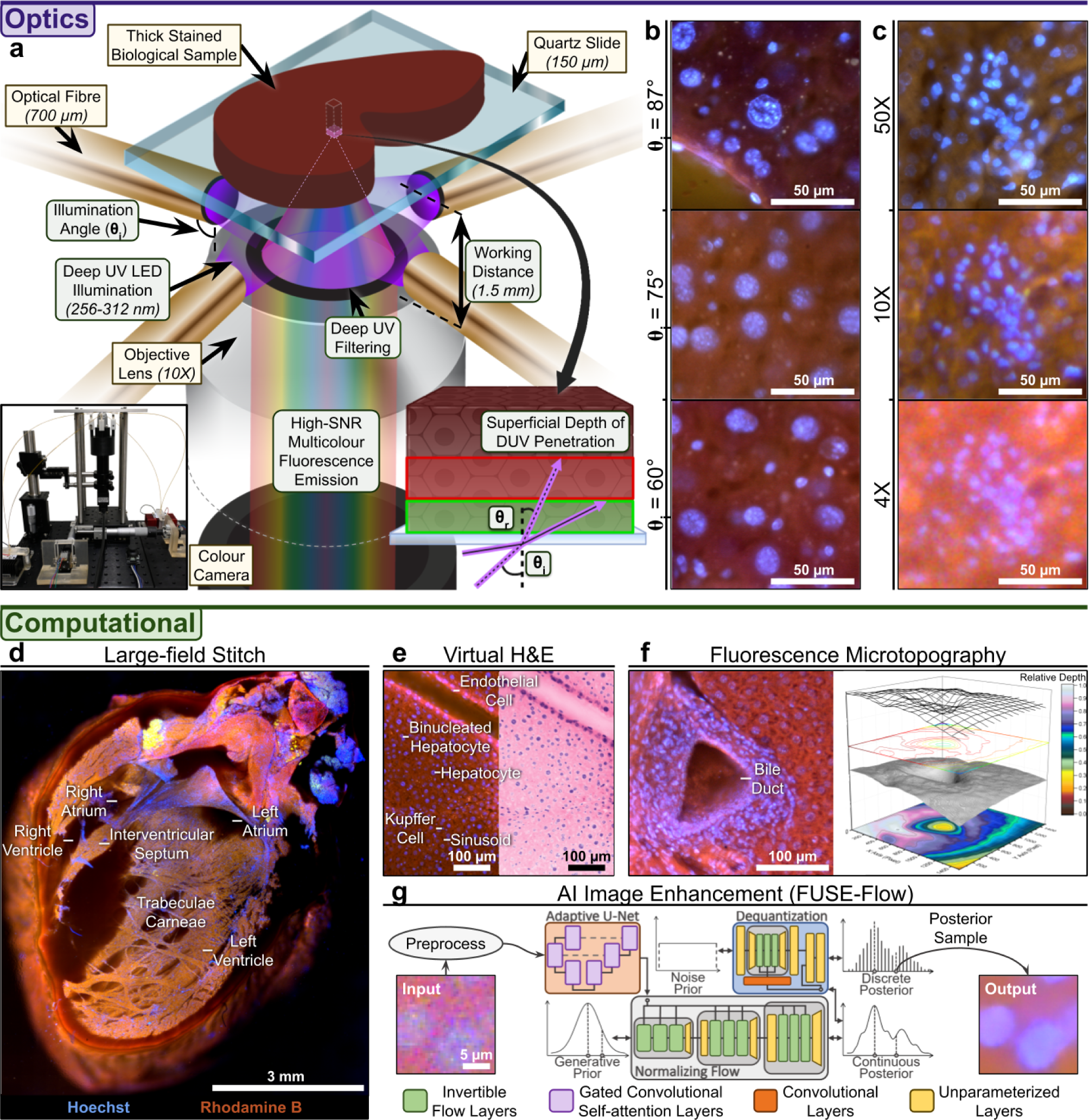
Optical and computational capabilities of Fibre optic microscopy with Ultraviolet Surface Excitation (FUSE). **a.** DUV illumination leveraged fibre optics for enhanced axial sectioning. **b.** Images of *>*1 mm-thick formalin-fixed mouse liver slice showed improved optical sectioning with increasing illumination angle at 50X. **c.** Multi-scale (4X, 0.10 NA; 10X, 0.25 NA; 50X, 0.55 NA) imaging of *>*1 mm-thick fresh mouse kidney slice showed nuclei-dense renal corpuscle. **d.** Stitched image depicted hand-cut cross-section of fresh rat heart, highlighting capability to quickly (*<*1.5 min) capture large (8 mm x 8 mm) regions of interest with facile (*<*2 min) sample preparation. **e.** Virtual H&E recolouring applied to fluorescence histological images to suit pathological workflows. **f.** Switching illumination between multiple optical fibres produced 3D fluorescence microtopography of mouse liver bile duct for textural imaging usable in advanced histological techniques. **g.** Conditional normalising flow enhanced low magnification (4X) images through learned statistical relationship between high-resolution detail and coarser structural elements.

Generative visual deep learning has greatly influenced the biomedical AI landscape following the impressive performance of generative adversarial networks (GANs) [20, 21] in image reconstruction and processing. The advent of diffusion [22] expanded the applicability of generative AI, now encompassing areas such as medical imaging [23] and computational microscopy [24, 25]. Normalising flows, a class of generative models less common in biomedical research, are popular in the physical sciences [26, 27] for their strong statistical basis. Unlike typical generative models that learn using approximations (VAEs [28] and diffusion models [29]) or adversarial mechanisms (GANs [30]), normalising flows precisely map complex non-parametric data distributions to familiar, tractable priors, such as the Gaussian or uniform distributions. Normalising flows consist of invertible layers, designed for computational efficiency and tractability, facilitating training through exact maximum likelihood estimation (MLE) [31–34]. Flow is potentially ideal when inferring conditional data distributions, a key aspect frequently sought in dependable biomedical AI applications. We built FUSE-Flow, a conditional normalising flow model that learns the statistical relationship between high-resolution detail and coarser structural elements (Fig. 1g), bringing variational inference, computational super-resolution and thick sample microscopy together for the first time.

A battery of studies was conducted to validate the capacity of this platform for biomedical applications at various magnifications (4X, 10X, and 50X). This included multi-scale fluorescence histology with 2-colour tissue staining protocols validated on standard H&E, immunofluorescence for protein expression quantification, quantitative cytotoxicity assays for organ viability, and subnuclear DNA statistics all on murine organ slices up to 0.4×8×12 *mm* and 1.3 million cells per surface.

## 2 Results

### 2.1 Fibre optic DUV illumination

Previous MUSE microscope designs had LEDs pointed directly at the sample, some with additional light guides or optics [14, 35]. While this approach delivers maximal optical power to the sample in principle, there is very limited control of the oblique angle of illumination due to practical considerations including the LED size and illumination cone, distance of the sample from the LED, and the optomechanical options available. While a previous study attempted to optimise illumination angle by the use of immersion medium [13], there has been little else reported on the potential for enhancing MUSE capabilities with more degrees of directional control.

Our setup used multi-mode fibres coupled to LEDs and positioned at the sample with customised miniature optomechanics. The flexibility and scalability of fibre optic illumination enabled fibres to be precisely adjusted in the polar axis (angle from horizontal), and multiple fibre sources to be positioned surrounding the sample at different azimuthal angles in the horizontal plane. While the compactness and arbitrary length of optic fibres provided logistical advantages such as the use of generic short-working-distance (down to *∼*1.5mm) objectives at 4X and 10X, fibre optic illumination further enabled two transformative advances.

First, illuminating at a near-horizontal polar angle produced optimal axial sectioning to the extent of enabling 50X magnification imaging. The use of higher magnification objectives enhances lateral resolution, but axial resolution (defined by the penetration depth of the DUV light in MUSE) also has a strong effect on image quality from a thick scattering sample. While angle here refers to the illumination direction of the source, light in fact emits from an optical fibre or LED as a cone, of which the marginal rays would travel deeper into the sample. Near-horizontal illumination ensured that even the marginal rays of the light cone would travel minimally in the axial direction. Our experiments showed that oblique illumination at unconstrained large angles produced blurred images even at 50X magnification. Only when the illumination angle was near-horizontal could cellular images of subnuclear resolution be obtained. It would be extremely inconvenient, if not impossible, to achieve such an illumination angle with direct illumination from a bulky LED even with focusing.

Second, optical fibres enabled sequential switching of single illumination directions to generate a set of images that could be used to produce a 3D reconstruction of the sample’s microscopic surface (Fig. 1f; Supp. Fig. 1). Obliquely illuminated images, while 2-dimensional, often showed a quasi-3D effect on uneven surfaces due to texture and shadowing [12]. We used the photometric stereo algorithm [17] to estimate depth from a single fixed field-of-view using fluorescence 2D images excited from multiple azimuthal directions. The near-horizontal illumination produced strong shadowing that highlighted not only prominent microstructural features such as tubules/ducts but also more minute textural differences that could be associated with tissue/cellular heterogeneity but would be indiscernible from thin-sectioned tissue or brightfield microscopy.

### 2.2 Image enhancement with conditional normalising flows

Accurate visualisation of nuclei is crucial for numerous pathological applications, particularly in cancer assessment. Greater magnifications are ideal for such applications as low magnification objectives often produce images with noise and poor colour contrast, hindering effective nuclear characterisation. However, higher magnification objectives introduce complex technical challenges, such as increased illumination requirements, limited fields-of-view, shortened working distances, diminished depths-of-field, and more pronounced optical aberrations. Typical solutions necessitate added hardware complexity, which precludes affordability and simple usage. Additionally, the ability to infer and convey uncertainty is a sought-after feature in contemporary biomedical AI applications [36]. We developed FUSE-Flow to enhance the image quality from a 4X objective for precise nuclear margin delineation, maintaining the valuable advantages of low-resolution objectives while achieving high optical clarity and communicating uncertainty.

The training of a model for our unique fresh tissue preparations and general-purpose application presented a multi-faceted challenge. This complexity prevented the use of common training strategies for image super-resolution such as supervision [37] (using perfectly registered high and low magnification images as targets and inputs respectively) and self-supervision [38] (where the inputs are modified low-resolution versions of the high-resolution targets). Obtaining perfectly registered targets and inputs was challenging due to the inherent dynamic nature of fresh tissue, largely credited to the presence of moisture that can induce real-time fluidic shifts, altering tissue morphology and positions. Conversely, non-supervised augmentation of the target to match the input domain is not viable, given the difficulty in precisely emulating the photometric distortions specific to each optical and experimental setup. These variations arise from factors such as differing chromatic characteristics of magnification objectives, fluorophore photobleaching, sample degradation, human error in preparation, and fluctuations in environmental conditions. We developed a unique semi-supervised strategy for the training of FUSE-Flow tailored to our distinctive fresh tissue imaging application. This approach was assessed on a common benchmark image dataset CelebA [39] (Supp. Fig. 9) as well as fluorescence histological images of sliced fresh mouse kidneys and acquired using FUSE. The model was trained using 10X magnification images as targets, while the inputs were augmented versions of these targets, with supervised adjustment to match the style (colour and resolution) of the corresponding loosely registered 4X magnification images. Subsequently, we evaluated it on a held-out (not part of the training dataset) kidney slice imaged at 4X magnification, with corresponding 10X images as references for comparison.

FUSE-Flow, along with a preprocessing step, effectively resolved nuclear boundaries with high contrast to adjacent cytoplasm, mirroring that observed in the 10X references (Fig. 2a,b). Generated images retained coarse structural features such as the position and general shape of nuclei, macro tissue architectures, illumination uniformity (Fig. 2c), and consistent focal clarity (Fig. 2d). Higher magnifications, especially with greater NA, exacerbate non-uniform illumination due to the increased light intensity demands. Despite the use of four optical fibres, the illumination from each fibre retains its intrinsic radial illumination pattern with a central peak intensity, making perfectly uniform lighting particularly challenging. Additionally, depth-of-field diminishes substantially at increased magnifications. Achieving distinct nuclear boundaries without these associated challenges, coupled with additional advantages such as broader fields-of-view and accelerated large-field imaging rates, bolster the platform’s capability.

**Fig. 2:**
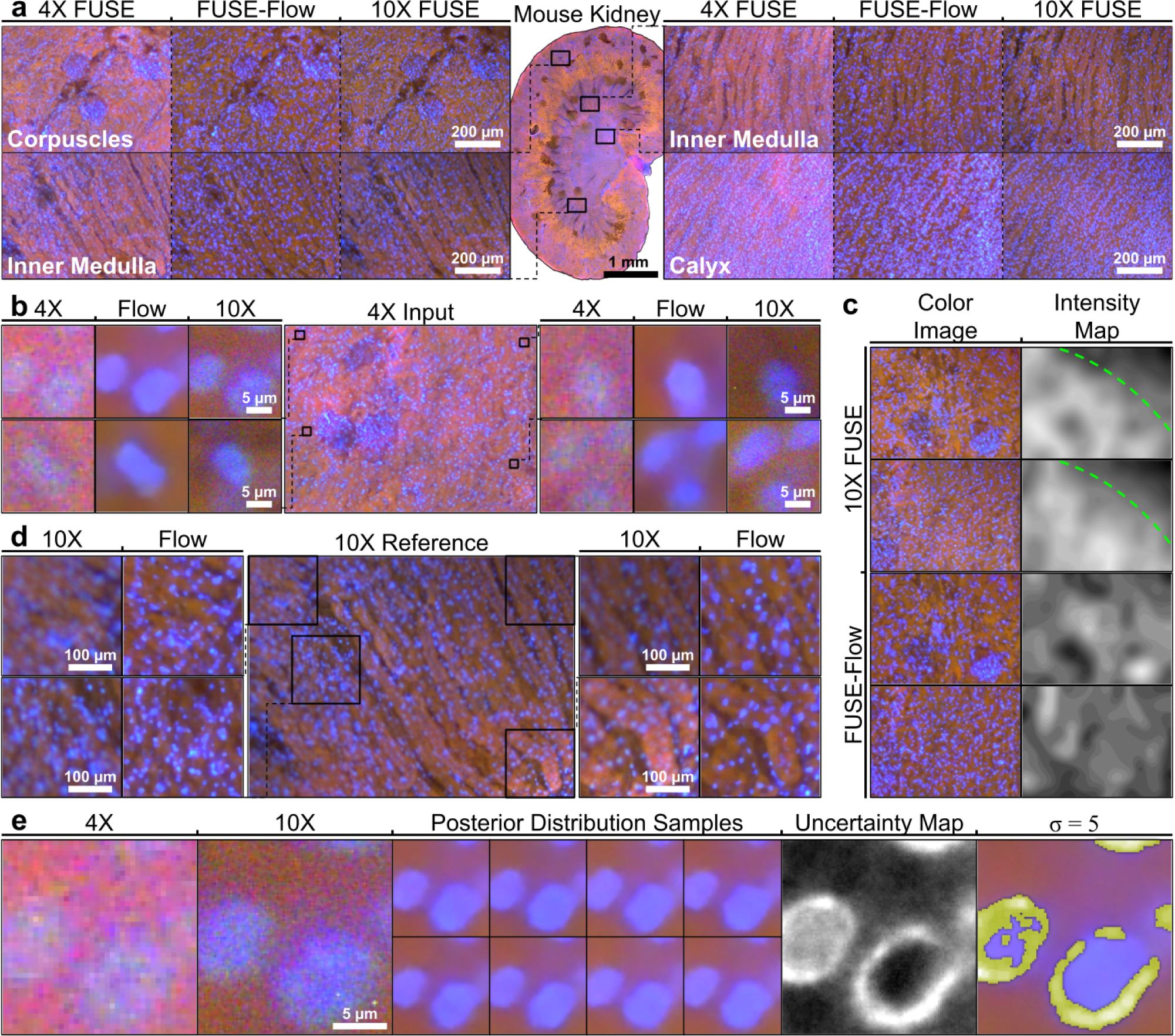
Image enhancement using FUSE-Flow. **a.** Performance overview on fluorescent histological images (held-out) of fresh mouse kidney slice. FUSE-Flow performed domain alignment of input 4X images to reference images in colour and detail while preserving input’s coarser features like nuclei positioning and tissue texture. **b.** FUSE-Flow enhanced nuclear margin sharpness and increased contrast between nuclei and cytoplasm. **c.** 10X images displayed clear bias in upper right corner due to non-uniform illumination. Bias was absent in model-enhanced images as evidenced by intensity maps correctly corresponding to tissue features. **d.** FUSE-Flow outputs show no out-of-focus areas, typically seen in higher-magnification images due to tissue regions falling outside objective depth-of-field. **e.** Multiple samples (*n* = 64) drawn from learnt posterior distribution could estimate conditional standard error to identify regions with highly aleatoric uncertainty. *σ* = 5 (or p-value= 3*e^−^*^7^) was highlighted.

Ideally, we aim to deduce the conditional standard error of predictions to convey the confidence (or lack thereof) we should have in the predictions, as a metric of uncertainty. Given the exact nature of the posterior distribution learnt by the normalising flow model is unknown, direct inference using parametric families would be inaccurate. We employed Monte Carlo simulations to calculate pixel-wise standard deviation as an empirical estimation for standard error (Fig. 2e). Regions with a five-sigma (*p <* 3*e^−^*^7^; commonly used in the physical sciences to denote highly unlikely events) level of uncertainty highlight the aleatoric uncertainty inherent in this ill-posed challenge—low-resolution and noisy inputs lack the comprehensive information needed for definitive output generation.

### 2.3 Fluorescence histology of thick tissue

UV-excited fluorescence has a unique capacity to deliver nuclear-level functional insights from thick tissue samples with minimal complexity. It involves straightforward sample preparation, imposes no laser requirements, and is compatible with scientific RGB cameras, which readily support multiplexed imaging and can be processed with standard image analysis and spectral unmixing techniques. We found DUV excitation well-suited for a wide range of fluorescence assays, further facilitating multiplexed imaging with the colour camera (Fig. 3a). We demonstrated the use of Hoechst 33342 as an effective nuclear stain with Rhodamine B or Eosin Y as a counterstain for DUV-excited fluorescence imaging of tissue samples, serving well as an analogue for standard histology. SYTO 9 and propidium iodide (PI) were also investigated as a unique approach to high-throughput tissue histology, leveraging large-field viability readouts to complement structural insights based on nuclear density. While this stain combination is marketed primarily as a bacterial assay, these dyes were also documented to work on eukaryotic cells. Alexa Fluor labels typically conjugated to antibodies, such as AF488 and AF594, were also DUV-excitable and could be readily used for the visualisation of protein expression.

**Fig. 3:**
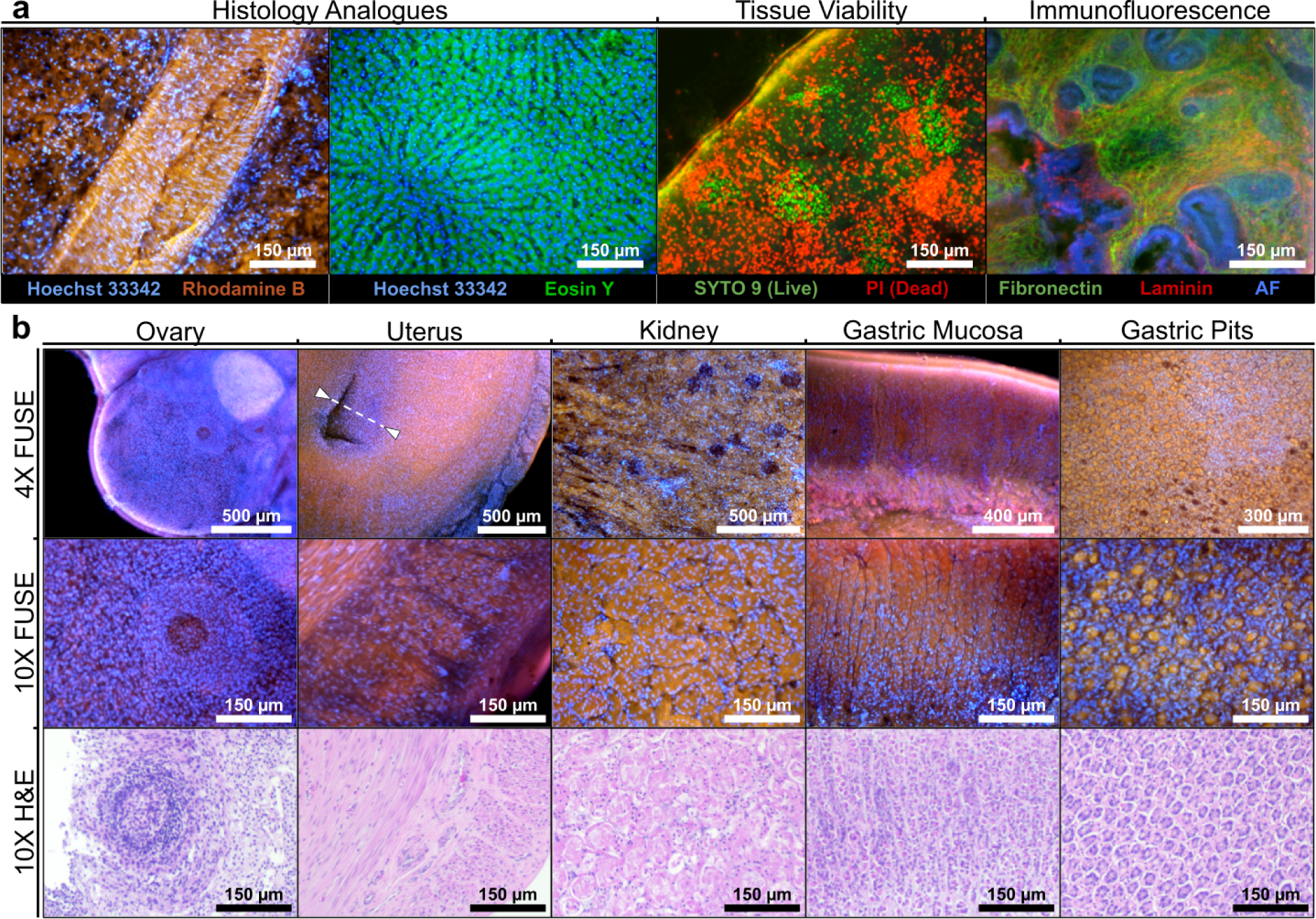
Fluorescence histology of thick fresh tissue. **a.** Range of stains that fluoresced upon DUV excitation. Nuclear (Hoechst 333422, SYTO 9, PI) and cytoplasmic stains (Rhodamine B, Eosin Y) served as analogues to H&E, revealing the inner surface of a blood vessel from fresh mouse kidney (Hoechst, Rhodamine) and surface of fresh rat liver (Hoechst, Eosin). LIVE/DEAD staining provided viability readouts of fresh renal tissue (SYTO9, PI). Immunofluorescence staining provided organ-level protein expression in fixed mouse uterus (Fibronectin, Alexa Fluor 488; Laminin, Alexa Fluor 594, AF: autofluorescence) **b.** Multi-scale imaging of Hoechst and Rhodamine stained fresh murine organs with H&E comparisons. Developing secondary follicle of the ovary was staged by the diameter of the follicle (white dashed curve). Stratum functionalis (white arrowheads and dashed line) was useful for staging of estrus cycle.

Prior studies had shown close correspondence between fluorescence histology and H&E histology [3, 12]. We broadly showcased this capability on our platform across the renal, digestive, and female reproductive systems (Fig. 3b). Fresh samples, which are important in clinical applications and biological viability studies, were our primary focus although we noted that fixed samples could achieve clearer preservation of cellular details and an even closer correspondence to histology. Tissue preparation protocols for DUV fluorescence histology did not appear to affect any downstream H&E processing, and the fluorescence images showed excellent nuclear visualisation with a high level of structural correspondence to the matching H&E images.

Large field-of-view coverage at cellular resolution enabled quantitative measurements at the organ scale that could correlate with physiology or phenotype, such as the size of a follicle indicating maturation and endometrial thickness as a marker of estrous stage. Quantitative FUSE analysis enabled the measurement of vascular structure dimensions and functional assessment of the uterus. Oblique illumination provided a shadow effect and a quasi-3D appearance, a unique perspective of the organ revealing the inner surface of the endometrium. The layer of simple columnar epithelial cells lining the endometrium corresponded well to histology. In rat kidneys, renal corpuscles, including the Bowman’s capsule and glomerular vasculature, could be distinguished, potentially revealing insights into kidney physiology and injury. High-quality *en face* views of the outer and inner stomach surfaces showed gastric pits and longitudinal muscle, respectively. While the visualisation of intact organs is limited to the surface, alternate imaging planes may be developed either by manual cutting or microtome. The cross-sectional structure of the gastric mucosa was studied via a ‘Swiss roll’ preparation of tissue cut, rolled up and mounted on its side.

Image-based quantitative estimation on our platform provided precise measurements while maintaining spatial information at cellular resolution. This technique was applicable to diverse biological contexts, ranging from single cells and 3D cultures (Supp. Fig. 8) to *ex vivo* tissue, with the caveat that information from deeper than the superficial surface could not be obtained. We applied this graphical quantitative approach to quantitative measurement in the various pilot investigations described in the following section.

### 2.4 Scale-specific pilot investigations

We demonstrated our platform’s versatility across key histological applications—two-colour H&E analogues, immunofluorescence, cell viability, and nuclear DNA content—through a series of hypothesis-driven pilot studies described in the following sections.

#### 2.4.1 Large-field viability analysis for renal toxicity from antibiotic insult

Vancomycin remains a standard treatment of methicillin-resistant *Staphylococcus aureus* (MRSA) [40] despite being linked to nephrotoxicity and acute kidney injury for decades [41, 42]. It is known to affect the proximal convoluted tubules of the kidney nephron due to the drug’s mechanism of action. We hypothesised that cell viability measurements at the organ scale could be a surrogate quantitative metric for toxic insults to renal function. Previous viability studies focused on cellular changes and lacked whole-tissue and regional analyses. Whole-tissue and regional analyses could potentially offer insight into the tubular specificity of vancomycin.

The SYTO 9 and PI viability assay based on membrane permeability visualised the effects of increasing vancomycin dosage, including standard treatment dosage [43, 44] and PBS control, at 4X magnification. FUSE imaging displayed diverse structures, revealing the kidney’s innate anatomical variability (Fig. 4a). Functional components like the renal calyx, corpuscles, and tubules were identifiable due to their unique nuclear density and response to vancomycin (Fig. 4b). Beyond visual comparisons, computer vision techniques enabled statistical insights. Pixel-wise comparison of the green and red colour channels respectively segmented live and dead cells, independent of brightness variation biases. To facilitate site-specific analysis, several cropping strategies were utilised depending on the specificity of boundary and difficulty of automation (Fig. 4c). Statistical evaluations showed significant (*p <* 0.0001) differences in viability between treatments and control groups in all regions (Fig. 4d). Notably, the medulla and cortex exhibited comparable viability patterns, indicating similar vancomycin effects. The corpuscle and tubule, however, manifested distinct viability distributions. Untreated corpuscles exhibited varied viabilities, indicating their natural predisposition towards cell death. Upon vancomycin exposure, most corpuscles exhibited drastically reduced viability. In contrast, the tubules’ viability distribution somewhat resembled that of the medulla and cortex but with much broader deviations, indicating a diverse treatment response. We observed a marked difference (*p <* 0.05) in viability between corpuscles and the surrounding cortex tissue across treatments (Fig. 4e), emphasising the heightened sensitivity and visual identifiability of corpuscles. This contrast was absent in tubules, despite known vancomycin-induced nephrotoxicity, and may be observable only from *in vivo* clearance. Although the clinical exposure of vancomycin to kidneys occurs during drug clearance in renal excretion and differs from that of our *ex vivo* kidney treatment, investigating the effects of antibiotic treatment on specific functional regions within tissue could still offer meaningful insight into the drug’s effects, while also relevant to more sophisticated *in vivo* tissue models. Our findings, influenced by individual variations and limited replicates, caution direct dosage comparisons. Nevertheless, our approach showcases the value of FUSE for drug evaluation in thick tissue models, bridging the gap between cell studies and clinical trials.

**Fig. 4:**
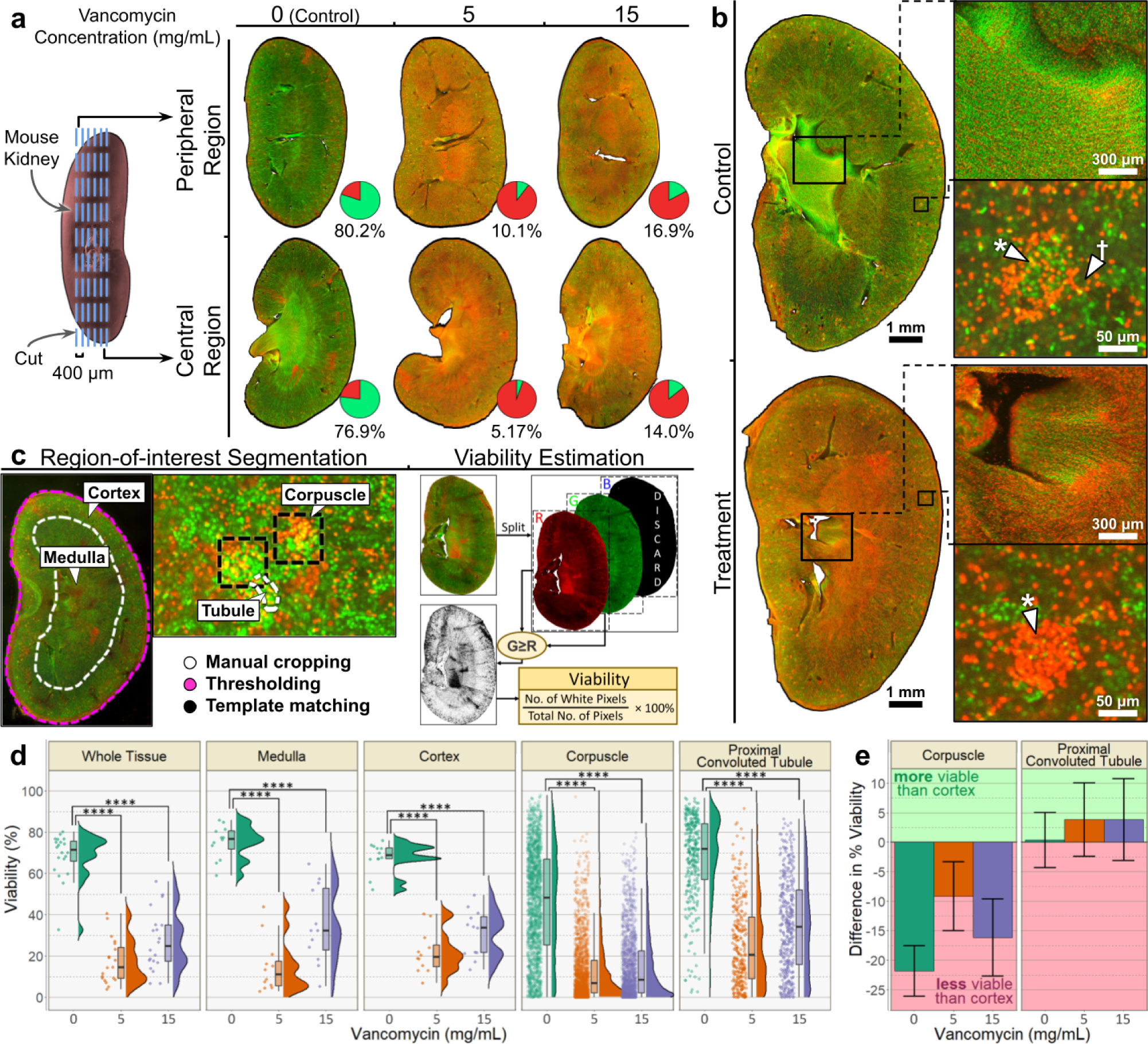
*Ex vivo* vancomycin antibiotic treatment on kidney tissue viability. **a.** Kidney slices from treated and control groups showed effects of vancomycin on various organ regions at 4X. **b.** Structure and stain intensity of renal calyx stood out from surrounding inner medulla. Renal corpuscles (marked by *∗*) were more susceptible to vancomycin toxicity. Decreased viability extended to proximal convoluted tubule (marked by *†*) of kidney nephron. **c.** Multiple segmentation strategies were employed to isolate specific regions of interest for downstream comparative assessment. Cells were deemed as live or dead depending on dominant image colour channel. **d.** Decline in viability highlighted toxicity of vancomycin. Distributional comparisons of regional viability estimates showed statistically significant (*p <* 0.0001) differences between treatment and control groups regardless of region. **e.** Emphasised contrast in viability of corpuscles and proximal convoluted tubules to their surrounding tissue (cortex), indicating consistent susceptibility to treatment. Error bars denoted confidence interval based on t-statistic calculated at 95% confidence level.

#### 2.4.2 Differential uterine protein expression quantification with immunofluorescence

The human female reproductive system displays early signs of ageing, marked by reduced fertility and hormonal dysfunctions [45]. While the mechanisms remain elusive, research indicates a correlation between extracellular matrix (ECM) protein composition and ageing [46, 47]. Fibronectin is a multi-functional protein that is important for embryogenesis, cell adhesion, and cell growth [48] that plays a major role in ECM assembly by binding to collagen, integrins, and other proteins [49]. Understanding the changes in protein composition that occur with ageing could provide insights into reproductive health and potentially help prevent the onset of diseases. In particular, fibronectin protein expression was found to be downregulated in the ovaries with ageing, impacting the ECM makeup and potentially leading to tissue degeneration [45]. Although ovarian dysfunction is the predominant factor impacting gestational outcomes, changes in uterine architecture may also be linked to decreased fertility. The relationship between uterine fibronectin expression and pregnancy has been investigated [50], but the effect of ageing on expression in the uterus remains elusive.

Immunofluorescence staining, followed by image-based quantification at 10X magnification, was employed to assess differences in uterine architecture and expression between mice at the onset (6 weeks) and end (32-36 weeks) of their reproductive life span (Fig. 5a). Expression level assesses antibody binding concentration, indicating protein abundance, while expression coverage reveals protein localisation. Aged mouse uterus slices exhibited statistically significant (*p <* 0.0001) differences in expression level from young uterus slices across all regions, indicating a decline in fibronectin protein expression with age (Fig. 5b,c). The distinct decrease in expression coverage further substantiated our hypothesis that fibronectin expression was downregulated with uterine senescence, raising questions about the potential impact on uterine functionality. Regional analyses indicated higher expression levels in the myometrium compared to that of the endometrium in both young (*p <* 0.05) and aged (*p <* 0.0001) uteruses. However, expression coverage did not exhibit this trend, with contradicting results in the young (n.s.) and aged uteruses (*p <* 0.01). Our study did not consider the estrous cycle of the mice, which may lead to differential expression patterns [51]. The quantification of protein expression may be sensitive to biases in illumination or other optical artefacts. Other protein analysis methods (e.g. western blot or mass spectrometry) may be required to substantiate any claims in specific regional expression differences. Limited information regarding differential expression patterns is known, but our preliminary study hints at similar patterns in fibronectin composition in uterine and ovarian senescence. The findings here highlight FUSE’s capacity for quantitative analysis of protein expression in extensive unsectioned tissue from relatively large organs.

**Fig. 5:**
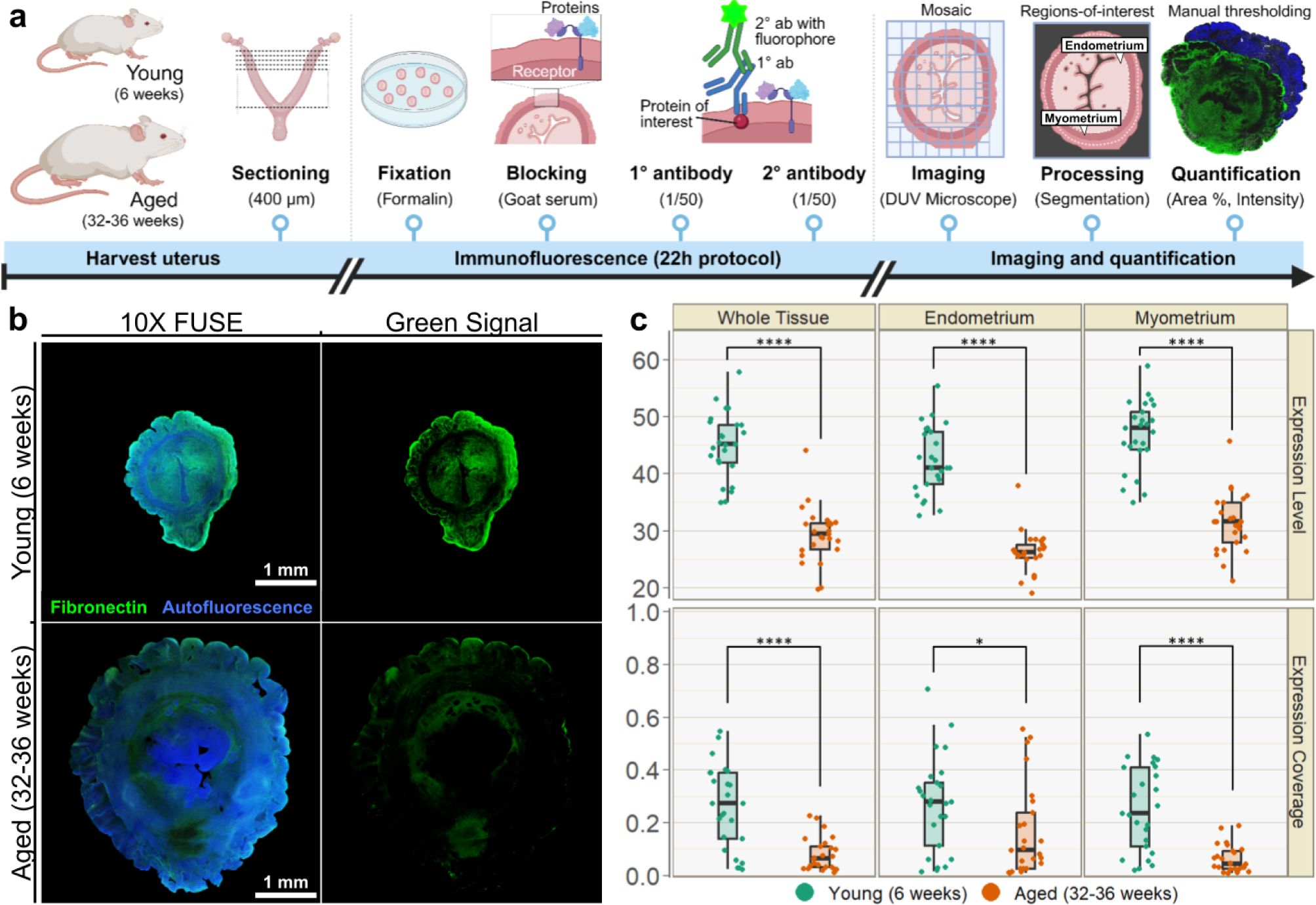
Uterine senescence on extracellular matrix fibronectin expression. **a.** Uterus slices (400 µmthick) from young (6 weeks) and aged (32-36 weeks) mice were subjected to anti-fibronectin immunofluorescence staining protocol prior to imaging and quantitative analysis. **b.** Increased fluorescence in young mouse uterus slices indicated higher fibronectin expression, correlating to well-understood changes in uterine architecture and reproductive health. **c.** Image-based statistical assessments (*n* = 25) revealed significant (*p <* 0.05) variations in fibronectin expression between the two groups. Expression level estimated fibronectin concentration from the signal intensity and proportion of tissue coverage reflected its distribution and localisation in tissue. No differential expression was observed between endometrium and myometrium, indicating a consistent decline in fibronectin with uterine ageing.

#### 2.4.3 Sub-nuclear characterisation of hepatic polyploidisation

Hepatocytes constitute the majority of the cellular makeup in the liver and its unlimited replicative potential is the key factor in the liver’s characteristic regenerative ability. They play a role in protein synthesis, carbohydrate and lipid metabolism, metabolic homeostasis, synthesis, storage, distribution, and detoxification of toxic compounds and regeneration [52]. Polyploidisation (duplication of chromosomes) of the hepatocytes is a unique feature of mature hepatocytes. It increases the DNA content of hepatocytes (diploid, 2n; tetraploid, 4n; octoploid, 8n; and so on) and has been linked to cellular senescence and stress (such as oxidative injury, surgical trauma, metabolic overload, and exposure to toxic insults). Many open questions regarding hepatocyte polyploidisation remain, such as the significance of the phases of hepatocyte polyploidisation observed in normal development and ageing, the role that altered hepatocyte polyploidisation plays in various pathological conditions, and the causal effect between rejuvenation of senescent hepatocytes and polyploidy reversal [53]. Obtaining nuclear-level information is critical to the study of hepatocyte polyploidisation and its significance during postnatal development and with senescence. In this investigation, we probe the potential of 50X FUSE to enable a simple, quick, and accurate quantification workflow of analysing DNA content and subnuclear detail of thick tissue samples.

Nuclear size distributions in hepatocytes sampled from liver slices of young (6 weeks) and aged (32-36 weeks) mice reveal three distinct trends (Fig. 6a). Firstly, the observed multimodal distributions align with recognised hepatic polyploidy phases [54]. Secondly, a right-shifted distribution of the aged population indicated a general increase in the DNA content of nuclei in aged mice. This is consistent with prior studies that found escalating hepatocyte DNA content with age, even after accounting for the polyploidy phase [55]. Lastly, the aged group displays a notable right skew, with its 2n frequency peak matching the 4n peak. In contrast, the young group’s 2n frequency peak is approximately twice its 4n peak, implying increased polyploidisation and proliferation in higher-order polyploids in the aged group. The hepatocyte cell cycle phase could be inferred from the sub-nuclear DNA content, as shown with H&E comparisons (Fig. 6b). Currently, obtaining such sub-nuclear insights requires thinly sectioned liver, single-cell cultures, fine needle aspiration, or complex and costly systems to optically section and provide sufficiently high axial resolution. While our findings are well supported by prior work, it should be noted that we did not account for any specific liver lobule and location from which the samples were harvested. Hepatocytes located at the origin (portal tract) may be younger than cells near the hepatic vein [55], and this relationship could affect the DNA content and nuclear area distributions within this study.

**Fig. 6:**
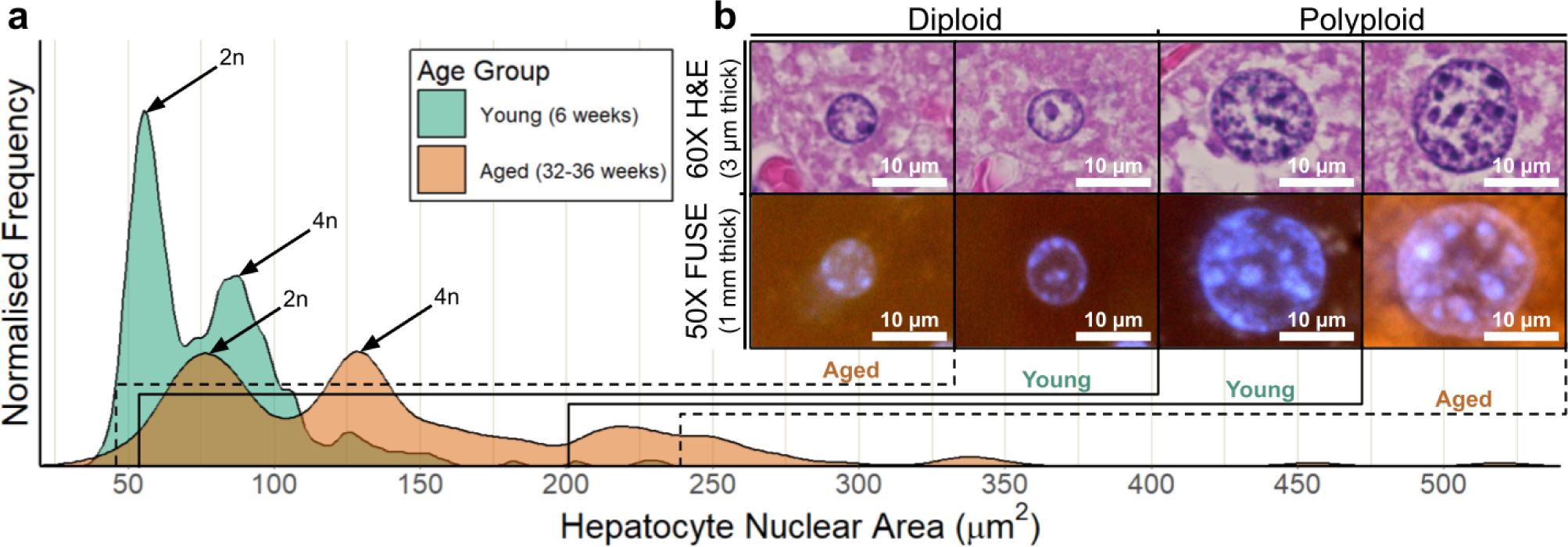
Senescence on hepatic polyploidisation. **a.** Multimodal distribution of hepatocyte nuclear area from young (6 weeks) and aged (32-36 weeks) mice with peaks corresponded to occurrence of hepatic polyploidisation (diploids, 2n; tetraploids, 4n). Rightward translation in distribution of aged population indicated a general increase in DNA contents of nuclei regardless of polyploidisation. Right skew of aged group suggested increased polyploidisation occurrence, with increased presence of higher-order polyploids in place of diploids. **b.** Fluorescence imaging (50X, 0.55 NA, air immersion objective) of diploid and polyploid hepatocyte nuclei alongside higher powered brightfield H&E images (60X, 1.40 NA, oil immersion objective). Similarly sized hepatocyte nuclei (not exact correspondence to FUSE images) from same tissue sample revealed close correspondence in sub-nuclear structures.

## 3 Discussion

DUV-excited fluorescence microscopy, also known as MUSE, has been celebrated for its simplicity and effectiveness in concept - the combination of oblique DUV illumination and widefield detection. However, prior published illustrations of real-world implementations [14, 15, 35, 56–58] showed inevitably large and complex setups, with the notable exception of Pocket MUSE, which is generally incompatible with large surgical or mesoscale samples [16]. Also notable in the MUSE literature was the conspicuous omission of imaging capabilities at *>*40X magnification, a mainstay of biomedical inspection and unusually difficult in MUSE due to the classically awkward implementation of oblique illumination. Our FUSE concept of fibre optic-delivered DUV illumination was not only space-efficient but also enabled a number of technical innovations that significantly expanded the capabilities of MUSE while still a respectful homage. Light delivery via a set of small diameter and easily micro-positioned fibre tips greatly relaxed the logistics of oblique illumination, to the extent of enabling the economical use of no-brand, short working distance objective lenses normally for standard widefield inspection. Fibre illumination at near-horizontal angles was also found to greatly enhance DUV optical sectioning for high-power microscopy at 50X magnification, which for the first time inducts MUSE into the family of multi-scale modalities. The fibres used in our work had to be relatively large in diameter (600 *µm*) to capture sufficient light from the LED. Fibres at this diameter still retain some rigidity, which may or may not be preferred when designing practical embodiments. Higher coupling efficiencies into smaller aperture fibres will eventually require laser sources and coupling optics.

Researchers have appreciated the phenomenon of meaningful shadowing produced by oblique DUV illumination, which generated a pseudo-3D effect from coarsely-cut samples and organ surfaces [12]. While multi-directional illumination would already improve uniformity, we further exploited directional control to produce 3D surface micro-topography using photometric stereo [17]. Fluorescence is typically assumed to be Lambertian (isotropic) [59] and has been previously studied as a feasible modality for photometric stereo reconstruction [60]. Although we did not exploit micro-topography for a comprehensive study beyond technical validation, we believe the technique could have applications in rapidly evaluating the morphology of 3D models as they gradually form microstructure or texture, which may reflect cellular heterogeneity and could have future relevance as a malignancy marker in histopathological evaluation [61], or even highlighting paths of low resistance for cell migration [62].

Generative AI has electrified the microscopy field, with extremely high-fidelity image reconstructions for resolution enhancement or modality transfer enabled by adversarial models [63]. However, there are several issues inherent to the adversarial training strategy of classical GANs [64], such as the lack of a defined convergence state, vanishing gradients [65], and mode collapse [30]. These issues have tempered the enthusiasm for GANs to be more broadly used in biomedical microscopy, where most granular image observations may be used to prove a hypothesis or make a diagnosis. The widespread acclaim for visual generative AI in the year 2023 could be arguably attributed at least in part to the rise of variational models, which include diffusion, variational autoencoders and flow. We chose flow purely because it is a classic technique underused in bioimaging for the direct statistical modelling of the data distribution, and not necessarily in favour of diffusion, which learns a denoising process and has similar properties. In demonstrating variational image generation in use cases where mode collapse or hallucination would be catastrophic, we argue that this class of models are particularly critical for principled yet high-quality generative imaging in biomedicine and microscopy. FUSE-Flow reconstruction of 10X data from 4X magnification has great practical value since 10X is the predominant setting used in MUSE, but it also provides a codebase and scalable framework for even more ambitious efforts in computational super-resolution.

Although multiple studies have already shown traditional MUSE to generate exquisite images from unsectioned tissue potentially for intra-operative histopathology, our work also featured a selection of murine tissue examples loosely themed around relatively large organs. This was partly to demonstrate our unique optical design to be competitive with the state of the art, and also to present a few new perspectives based on large-field mapping to motivate the serious consideration of FUSE as a tool for quantitative tissue biology. While we acknowledge that fixation is ideal for visualisation, we presented several fresh and unfixed biological preparations to envision a translational step towards DUV excitation for live samples and *in vivo* models, both important use cases in biological microscopy. Real-time live imaging could even have potential clinical applications in surgery including anastomotic viability and margin evaluation. Despite the well-known mutagenic effects of DUV light, the very brief sub-second exposures needed for high-quality imaging may be tolerable, and motivate dosimetry studies. Given concurrent advances in DUV light sources, fibre optics and microscopy, the potential of minimally invasive ultraviolet endoscopy [66] for real-time cellular interrogation of living organs could be due for a revisit.

The keystone of our multi-disciplinary effort is a curated set of hypothesis-driven biological investigations that are modestly scoped to corroborate known science, namely antibiotic toxicity, ECM expression, and polyploidy while demonstrating a novel quantitative approach using thick-sliced tissue sampled from a relatively large set of intact organ samples. While we emphasise the ability to infer histological and biological insights in a large region of interest without the need for thin sectioning, being constrained to surface-level information can be restricting for 3D biological samples; deeper information beyond surface texture is largely unobtainable. FUSE presents a new opportunity for biologists to rapidly interrogate coarsely sliced tissue at the cellular level from numerous candidates that would normally be logistically overwhelming to work through with standard confocal microscopy or cleared light-sheet microscopy. This high-throughput approach could work as a screening modality much like a tissue microarray at ‘mesoscale’, for which the most promising candidates can be selected for more thorough standard investigation. As the science of tissue omics accelerates and the lines between clinical pathology and tissue biology continue to blur, we humbly anticipate FUSE and FUSE-Flow to represent important inroads made towards a fast platform technology for multiplexed, high-throughput tissue insights.

## 4 Methods

### 4.1 Microscope design and components

The deep ultraviolet (DUV) fibre optic fluorescence microscope platform was based on an epi-illuminated inverted microscope design. The microscope sat on a 30×30 *cm* optical breadboard and comprised the illumination, optical, sample holder, and mechanical and electrical components (Supp. Fig. 2a).

#### 4.1.1 Illumination components

DUV illumination at a peak wavelength of 277 *nm* was produced by DUV LEDs (ILR-OV01-O275-LS030-SC201.; Osram) powered by a constant 700 *mA* current supply (RECOM) and cooled with a heat sink. Magnetic butt-coupling of multi-mode optical fibres (57-070; Edmund Optics), with an outer diameter of 710 *µm*, to the light-emitting diodes (LEDs) was achieved with a custom 3D-printed magnetic fibre coupler (Supp. Fig. 2b). The magnetic coupler allowed hot swapping of the illumination source and the optical fibres. The butt-coupled optical fibres had a measured output power of around 6.29 *mW* and a coupling efficiency of roughly 5%. The DUV-emitting end of the fibre optic was controlled via a 3D-printed custom optomechanical fibre holder jig which provided four degrees of freedom, allowing fine control and angle adjustments of the fibre for uniform sample illumination. An extreme oblique incident angle of illumination (*∼* 87°) was used in the 50X objective setup and (*∼* 85°) in the 10X objective setup. The incident angle of illumination was (*∼* 60°) for the 4X objective setup (Supp. Fig. 2c). With the 4X, 0.1 NA objective, sharp micrographs could be obtained without requiring the fibre optic to be positioned at a near-horizontal angle (Supp. Fig. 5). The smaller incident angle (*∼* 60°) of illumination was preferred for 4X as it provided stronger illumination while still maintaining sharp images. The area of illumination for the 10X and the 4X setup was measured to be approximately 4.88 *mm*^2^ and 8.58 *mm*^2^ respectively. The intensity of the 10X and the 4X setup was calculated to be approximately 1.29 *mW/mm*^2^ and 0.733 *mW/mm*^2^ respectively. The area of illumination and intensity of the 50X setup is estimated to be similar to the 10X setup.

#### 4.1.2 Optical components

Digital images were taken using a colour camera (BFS-U3-120S4C-CS; Teledyne FLIR) magnified by either 50X finite conjugate telecentric objective (375-052; Mitutoyo), or 10X or 4X finite conjugate DIN objectives (43-907, 67-706; Edmund Optics). The working-distances for the 50X; 0.55 NA, 10X; 0.25 NA, and 4X; 0.10 NA objectives were 13.00 *mm*, 1.50 *mm* and 15.25 *mm* respectively. A custom 3D-printed objective holder was non-mechanically coupled to a tube assembly (54-868; Edmund Optics), enabling hot swapping between the 50X, 10X, and 4X objective setups (Supp. Fig. 2c). The objective holder included a filter slot that enabled suppression of specific wavelengths, which could minimise unwanted signals, including autofluorescence from samples. The tube assembly connected the objective holder with the colour camera. The effective resolution was measured by conducting the knife-edge resolution test across the sharp edge of a blade. The 10 - 90% intensity transition, which corresponds to the Rayleigh resolution, was measured at 9 different points and averaged to obtain the half-pitch spatial resolution. The half-pitch spatial resolutions were measured to be 3.60 *µm*, 1.11 *µm*, and 0.311 *µm* for the 4X, 10X, and 50X objective setups respectively. The effective field-of-view was determined to be 0.748 by 0.561 *mm* (10X) and 1.85 by 1.39 *mm* (4X) by imaging a checkerboard calibration target. For 50X, the effective field-of-view was calculated to be 0.150 by 0.112 *mm*.

#### 4.1.3 Sample holder, mechanical and electrical components

Histology cassettes were modified to include a 150 *µm* thick UV-transmissible quartz window to contain tissue. A custom-machined aluminium sample holder supported by four optical posts was used to mount the histology cassettes. Standard histology cassettes (C33118001PH, Uni-Sci) were modified by cutting a hole approximately 11×20 *mm* before applying liquid epoxy to adhere a quartz coverslip (JGS2, Latech) to the outer surface covering the hole. The quartz coverslips acted as a flat and UV-transmissible surface for samples to be placed on for imaging.

X-axis and Y-axis movement, at sub-micrometre precision, of the sample relative to the camera was enabled by mounting the sample holder on a stack of two manual linear translation stages. Z-axis movement was enabled by a manual vertical translation stage. Two 3D-printed stepper motor holders and stepper motor couplers were designed to co-axially couple the stepper motors to the micrometre knob of the stage. A microcontroller controlled the stepper motor by either parsing a set of sequential instructions allowing for automated stage movement or inputs from an analogue stick for manual stage movement of the microscope in the X and Y direction. The flashing of the LEDs was controlled by the microcontroller during image acquisition, synchronised to the camera’s exposure time, to reduce unwanted DUV exposure to samples during stage movement of the automated imaging process.

### 4.2 Computational details

#### 4.2.1 Conditional normalising flow for image enhancement

##### Architecture

Our conditional normalising flow was developed with inspiration from existing open-source implementations (Supp. Fig. 3a). SRFlow [34], a recent demonstration of normalising flow for single-image super-resolution, formed the foundational structure for our model. We adopted advanced techniques from Flow++ [33], such as variational dequantization and the application of gated residual networks [67, 68], to enhance the model’s performance. Moreover, our design includes a custom-made adaptive U-Net tailored for arbitrary input sizes, ensuring the extraction of features with appropriate dimensions to feed the primary normalising flow. We also assessed the integration self-attention mechanisms, Squeeze-and-Excitation [69] and Convolutional Block Attention Module [70], tailored for convolutional neural networks. Ablation studies, performed on the CelebA [39] dataset, are detailed in the supplementary material (Supp. Fig. 9,10).

##### Data

Fresh kidneys from a sacrificed mouse were immediately prepared and imaged post-harvesting. One kidney was sliced longitudinally and the other transversely, yielding 30 slices in total. From the extensive set of over 4,000 images from large-field imaging on both 4X and 10X magnification, a small subset of 36 pairs of high-quality images (in focus and lacking imaging artefacts) that displayed important structural features were selected. A comprehensive data processing pipeline (Supp. Fig. 3b) generated input and target patches from the 36 image pairs for model training. Fast non-local means denoising [71] despeckled the original images, enabling the model to concentrate on structural variations without noise-induced biases. The high-resolution images from the 10X set were simulated as low-resolution through imputation downsampling and blurring. Histogram matching aligned the colour profile of these simulated images with the corresponding 4X images. The final processing step involved splitting the images into 96×96 *px* and 24×24 *px* patches (target and input respectively) for training, with selective minor blurring to mitigate any artefacts introduced from the previous step. The training utilized a set of 120,000 patch pairs.

##### Training

The model was trained on an NVIDIA RTX A6000 GPU (48 GB memory) over 48 hours, spanning 32 epochs with a batch size of 4. We set the learning rate at 1.0*e^−^*^4^, decaying by a multiple of 0.9 each epoch. The posterior was mapped to a standard normal distribution (Gaussian with a standard deviation of 1.0). The adaptive U-Net underwent pre-training with binary cross-entropy as the loss function, and once trained, it was frozen during the normalising flow training. During training, horizontal and vertical flips were applied with a probability of *p* = 0.5.

#### 4.2.2 Image acquisition and large-field mosaicing

##### Imaging settings and processing

Colour images were captured with exposure ranging between 0.1 to 3 seconds. The images were saved in PNG file format. Image processing was done with the open-source ImageJ (Fiji) [72] and limited to brightness and contrast, colour balance, and despeckling to remove hot pixel artefacts from the colour camera. No background correction, subtraction or deconvolution was performed on the images.

##### Automated image acquisition

Image acquisition was facilitated by an Arduino, Python code on a laptop attached to the microscope, and the Teledyne FLIR Spinnaker SDK. Manual image acquisition was done using an analogue stick with the Arduino for motor control and the SpinView application from the Spinnaker SDK for the viewing and saving of images. Automated image acquisition for large mosaic images was coordinated by our custom Python software. It received and processed user input, then controlled the camera through the Python interface from the Spinnaker SDK and the stage by sending condensed instructions to the Arduino.

##### Large-field mosaicing

Large-field mosaicing was enabled by the Python software integrated with the microscope and the freely available Microsoft Image Composite Editor. Images were automatically and sequentially captured with an adjustable overlap (usually set at 10%/15% margins) and stitched together using Image Composite Editor to achieve gigapixel-sized mosaic images comprising hundreds of individual images. The sample stage was precise enough for images to be manually focused prior to imaging, with the Python software subsequently automatically controlling the stepper motors movement, flashing of the DUV LEDs, and the camera capture of images, without additional refocusing required, over the entire sample area.

#### 4.2.3 3D microtopography with photometric stereo

##### Sequential illumination and image acquisition

To obtain the micro-topographical information over a region of interest on a sample, each of the four fibre optics was coupled to an independently controlled DUV LED. A reference image of the region of interest was then captured with illumination coming from all the fibre optics. Four sequential images with directional illumination from each fibre optics were obtained while maintaining the same field-of-view over the region of interest. The exposure of these four sequential images was increased (up to 3 seconds exposure) compared with the reference image, to compensate for the dimmer illumination coming from a single fibre optic.

##### Image processing

Despeckling was performed on all images using ImageJ to remove hot pixel artefacts as the bright pixels which would affect the subsequently obtained surface normals map. Following this, the four sequential images were histogram-matched with the reference image to obtain the different direction-of-illumination images with the same relative brightness. Histogram matching was done to force a more uniform brightness between the four images given there were slight variations in illumination intensity between each of the four optical fibres.

##### Photometric stereo for surface normals and depth map

With the four histogram-matched different direction-of-illumination images, four representative vectors for each direction of fibre optic illumination were then inputted into our implementation of the photometric stereo algorithm (vectorised and ported to Python, based on an open-source Matlab implementation [73]). The four representative vectors reflected the 90° rotation about the image’s centre to match our setup’s fibre orientation. The photometric stereo algorithm produced the surface normals and depth map of the region of interest (Supp. Fig. 1). It was noted that changing the input vector angle of illumination (obliqueness of illumination angle) did not significantly affect the obtained depth map as the depth map displayed the normalised depth rather than true depth. The depth map was rendered in 3D, with either the H&E recoloured image or the despeckled reference image used as the surface texture in PyVista, to produce an interactive 3D micro-topographical reconstruction of the sample region of interest.

### 4.3 Biological studies methodology and preparation

#### 4.3.1 Histological studies

For tissue samples, female NCr nude mice were euthanised at the humane or experimental endpoint of a separate study in accordance with the approved A*STAR Institutional Animal Care and Use Committee (IACUC) protocol. Mice were dissected and routine post-study tissue collection was performed. Alternatively, whole mice and rats carcasses or specific harvested organs were purchased (InVivos). Organs were harvested and washed in 1X Phosphate Buffered Saline (PBS) (BUF-2040-10×4L, 1st BASE), and sliced either using a vibratome (VF500-0Z, Precisionary Instruments) or manually using a scalpel depending on the use case. For vibratome slicing, the organ was embedded in 2% Type I-B agarose (Sigma Aldrich). In brief, the sample was glued onto the plunger of the specimen tube, liquid agarose was pipetted in excess to cover the tissue before the agarose was solidified using a chilling block. The specimen tube containing the sample was loaded onto the vibratome and sliced into 400 *µm* to 1 *mm*-thick slices at various oscillation and advance speeds depending on the organ type. The surrounding agarose was delicately removed using forceps before staining. Hoechst 33342 (62249, Thermofisher Scientific) and Rhodamine B (A5102, TCI) fluorescence stains were prepared at various concentrations for optimal staining of different tissue types. Tissue samples were stained for 30 seconds to 1 minute, before being washed in 1X PBS twice, for 30 seconds each. Samples were blotted dry with Kimwipes and placed on modified histology cassettes for imaging (Supp. Fig. 4). 0.1 *mg/mL* Hoechst with 2 *mg/mL* Rhodamine B was identified as an effective fluorescent concoction for general staining over a wide range of fresh and fixed murine tissue including organs like the heart, liver, stomach, and kidney, as well as the female reproductive tract. 0.1 *mg/mL* Hoechst with 4 *mg/mL* Rhodamine B was also used for higher contrast of structural features such as the tubules of fresh kidneys. Organs which featured a high level of nuclear concentration such as the ovaries and gastric surface of the stomach were better highlighted with a lower level of nuclear stain ratio of 0.1 *mg/mL* Hoechst with 1 *mg/mL* Rhodamine B and 0.05 *mg/mL* Hoechst with 2 *mg/mL* Rhodamine B respectively.

#### 4.3.2 Renal viability studies

##### Procedure

We harvested 12 kidneys from 6 mice and classified them into 3 treatment groups, control, low dose of vancomycin (5 mg/mL), and high dose of vancomycin (15 mg/mL). The fresh kidneys were sliced coronally into 400 *µm*-slices to obtain 5 mid-section slices per kidney, totalling 60 slices. Samples in the treatment group were treated with vancomycin (vancomycin hydrochloride, 1709007, US Pharmacopeia) for 1 hour. After treatment, kidney slices were washed in PBS before incubating in 5:1 SYTO 9 and PI nucleic acid stain (7.5 *µL* for 15 minutes with shaking (LIVE/DEAD BacLight Viability Kit, L7012, Invitrogen). Samples were blotted dry with Kimwipes and placed on modified histology cassettes for imaging.

##### Viability quantification

Our general strategy for image-based viability quantification involves 2 stages: segmentation followed by pixel-wise relative green and red comparison (Fig. 4c). Specific segmentation strategies differ for each use case. Colour comparisons begin with first splitting the colour image into its red, blue, and green (RGB) channels. A map of pixels where the green channel has a higher intensity than the red channel represents the live regions of the tissue is then created. This is a convenient strategy as the viability assay of choice stains in green (SYTO 9) and red (PI) at similar intensities. Since PI is only permeable to dead cells and has a stronger affinity for nucleic acids [74], the point where the red signal is greater than the green represents a significant uptake of PI and thus differentiates dead cells from live ones. This strategy is also robust to any local and global variation in brightness that tends to happen during the imaging process.

Single cells were segmented using Otsu’s method of thresholding before pixel-wise relative colour comparison. Kidneys were segmented from their background using constant thresholding. Kidney regions of interest like the cortex, medulla, corpuscles, and proximal convoluted tubules were cropped using various strategies. The cortex and medullas were manually cropped using ImageJ for 12 kidney slices per group with visible cortical and medullary regions. All discernible longitudinal proximal convoluted tubules were manually cropped from each kidney sample due to their non-uniform shape. Corpuscles were detected using template matching and cropped to a fixed size, generating over 4,777 crops. The Welch Two Sample t-test was used to determine the statistically significant differences between treatment groups (Fig. 4d,e). Statistical tests and plots were done using R programming language and Tidyverse [75].

#### 4.3.3 Uterine senescence studies

##### Procedure

We harvested 5 uteruses each from young (6 weeks) and aged (32-36 weeks) mice. Mouse uteruses were sliced transversely into 400 *µm*-slices. Mouse uterus cross-section tissue was fixed for 30 minutes in 4% formaldehyde (VWR) before blocking for 30 minutes in 10% Normal Goat Serum (50062Z, Life Technologies). Thereafter, primary antibody 1/50 Anti-Fibronectin (ab2413, Abcam) was added to the uterus slices and incubated for 16 hours. The slices were washed twice in 1X PBS before the secondary antibody Alexa Fluor 488 (A11034, Life Technologies) were applied and left to incubate for 5 hours. After incubation, samples were washed twice in 1X PBS prior to imaging. A total of 25 young uterus and 25 aged uterus slices were imaged at 10X magnification (Fig. 5a, created with BioRender.com.).

##### Quantification and analysis

Large-field mosaicing of the uterus images was enabled by Python code in post-acquisition processing. Gigapixel-sized mosaiced images were obtained. The image background and bright tissue borders in the perimetrium were removed by manual cropping using ImageJ. Uterus images were further cropped to obtain endometrium and myometrium areas for downstream protein expression quantification. Threshold value (*α* value) was ascertained for each individual tissue. *α* value was manually determined through analysis of pixels with prominent green intensity in its fluorescence signal. The selected *α* value returns a RGB image with adjusted blue intensity, and subsequently the blue B*>*G and G*>*B masks. Green mask was verified to be representative of the signal region. *α* values ranging from (0.3 to 0.7) were applied to select for the green signal with Alexa Fluor 488 tagged secondary antibodies binding, correlating to fibronectin protein expression in uterine tissue. Expression level was determined by the mean average green intensity of the image, with intensity values between 0-255. Expression coverage represented the expression area ratio of the green intensity over the tissue area. The Welch Two Sample t-test was used to determine the statistically significant differences between treatment groups (Fig. 5c). Statistical tests and plots were done using R programming language.

#### 4.3.4 Hepatic senescence studies

##### Procedure

For the young group, livers were harvested from 3 6-week-old mice, and for the aged group, livers were harvested from 3 32-36 weeks-old mice. The 6 livers were fixed overnight in 4% formaldehyde (VWR). Following overnight fixation, tissue preparation was largely similar to the fresh tissue sample preparation methodology described in 4.3.1. The 6 livers were then transferred to 1X PBS before being sliced to 1 *mm* with a vibratome. The fixed liver slices were stained for 6 minutes in a stoichiometric DNA labelling agent, Hoechst 33342 (0.4 *mg/mL*) and Rhodamine B (2 *mg/mL*) mix. The stained livers were placed on the modified histology cassettes and gently compressed with biopsy pads. After imaging, the stained livers were processed through the usual H&E workflow and imaged on a brightfield microscope (Nikon ECLIPSE Ni-E, DS-Ri 2) with a high powered PlanApo 60X, 1.40 NA oil immersion objective.

##### Image analysis

10 images were obtained from each liver using the 50X objective. From the 60 images (30 images from young mice, 30 from aged mice), hepatocyte nuclei were manually cropped using the elliptical tool in ImageJ (Fiji) [72]. 421 and 207 hepatocyte nuclei were cropped from the images of the young and aged mice livers respectively. The area of the hepatocyte nuclei was measured in ImageJ (Fiji). The hepatocyte nuclei area was plotted against the normalised frequency to obtain the frequency distribution of hepatocyte nuclei sizes for both the young and aged groups of mice.

## Acknowledgements

We would like to thank Dr Chee Bing Ong for his invaluable support and pathology-related consultations, Dr Nazihah Husna Abdul Aziz for her input into various experiment designs, Zhe Li Ha for her artistic contributions to figure design, Li Qin Shen for her contributions to software development for microscope automation, Stefanie Zi En Lim, Jeremy Rui Quan Lee for their contributions to data preparation, Rachel Yixuan Tan for her contributions to initial mechanical designs, and Dr Zesheng Zheng for engineering-related consultations. We would also like to acknowledge Nikon Imaging Centre (NIC) @ Singapore Bioimaging Consortium (SBIC) for permitting access to their confocal and brightfield imaging systems.

## Author contributions

Conceptualisation: KL, JLYA, ASKY, KHT, CJJT, RHKS

Methodology: ASKY, KHT, JLYA, KL, CJJT, RHKS, CWXT, JSJK

Investigation: KHT, ASKY, JLYA, CWXT, JSJK, KL, JYHT

Visualisation: JLYA, ASKY, KHT, CWXT, JSJK, JYHT

Supervision: KL

Writing-original draft: KL, ASKY, KHT, JLYA

Writing-review & editing: KL, JLYA, KHT, ASKY

## Data and materials availability

We open-sourced our PyTorch implementation of the conditional normalising flow model for image enhancement. It is available at https://github.com/KaichengGroup/FUSE-Flow. Code used for image processing and statistical analysis is available from the corresponding author upon reasonable request. All data needed to evaluate the conclusions in the paper are present in the paper and/or the Supplementary Materials. Experimental data underlying the results presented in this paper are available from the corresponding author upon reasonable request.

