## Supplementary Materials for "Multi-scale tissue fluorescence mapping with fibre optic ultraviolet excitation and generative modelling"

Tissue mapping with generative UV microscopy

Joel Lang Yi Ang[†1,3], Ko Hui Tan[†1], Alexander Si Kai Yong[†1],
Chiyo Wan Xuan Tan[4], Jessica Sze Jia Kng[4],
Cyrus Jia Jun Tan[4], Rachael Hui Kie Soh[4], Julian Yi Hong Tan[4],
Kaicheng Liang*[1,2,3,5]
**

[1]Institute of Molecular and Cell Biology, [2]Institute of Microelectronics,
[3]Bioinformatics Institute, [4]Institute of Bioengineering and Bioimaging,
Agency for Science, Technology and Research (A*STAR), Singapore
[5]Lee Kong Chian School of Medicine, Nanyang Technological University (NTU), Singapore

[†]Equal contribution, ordered alphabetically
*Corresponding Author

Oct 2023

**This PDF File includes:**

- Supplementary Text
- Supp. Fig. 1 to 10

### 1 Supporting Investigations

#### 1.1 Thick tissue imaging at varying magnifications with FUSE

We demonstrate the ability to obtain corresponding features at different magnification levels (4X/10X/50X) from unfixed thick tissue samples (Supp. Fig. 6). Low magnification imaging could serve as a means of high throughput screening of multiple samples at the organ level. Specific features or key regions of interest could then be identified and imaged for subsequent analysis at higher magnification and resolution. The use of magnetic butt coupling of the fibre optics to the DUV illumination source with the non-mechanical coupling of the objective holder to the focal tube allows hot-swapping between different objective setups (Supp. Fig. 2). With the 4X, 0.1 NA objective, sharp micrographs could be obtained without requiring the fibre optic to be positioned at a near-horizontal angle (Supp. Fig. 5). This is likely due to the depth of field of the low NA objective being larger than the depth of penetration of DUV in thick tissue samples. For the 10X and 50X objective setups, the fibre optic was angled such that the illumination was directed at a near-horizontal plane. This was shown to improve the image quality by reducing the depth of DUV penetration to more closely match the small depth of field of the high NA 50X objective. Other studies employing UV surface excitation have sought to reduce the depth of penetration through the use of a high refractive index immersion medium. While we demonstrate 50X FUSE with air immersion objectives and fibre optics to reduce the depth of DUV penetration, FUSE allows for an extreme horizontal angle of illumination that could be paired with a high refractive index immersion medium for even higher magnification objectives to be used.

#### 1.2 Quantitative measure of hepatocyte DNA content

**Cell cycle phase analysis through quantification of DNA content in thick samples** To demonstrate the ability to obtain subnuclear information from thick tissue samples with FUSE, we study the DNA contents within the hepatocyte nuclei from thick liver samples ($> 1\ mm$). The blue channel of high magnification (50X) images were Gaussian-filtered and then automatically thresholded using Otsu's method (Supp. Fig. 7). The number of blue pixels above Otsu's threshold and within the hepatocyte nucleus envelope was then divided by the total number of pixels of the hepatocyte nucleus to obtain the DNA area expression (%) within the hepatocyte nucleus envelope. This was performed on a pair of binucleated hepatocyte nuclei and a pair of similarly sized mononucleated hepatocyte nuclei as a means to obtain a simple yet quantitative comparison of a subnuclear (DNA content) feature, providing cell cycle phase information.

#### 1.3 Cell viability in cell cultures

Conventional viability measurements use fluorescent plate readers (spectrophotometers), which give fluorescence intensity readouts summed over entire samples but lack spatial information or cell-level statistics. We showcase an automated image-based fluorescence assay to determine cell viability (Supp. Fig. 8b). MCF-7 cells (stained with SYTO 9 and PI) were chemically insulted and 60 images were obtained every minute. An automated cell segmentation method using Otsu's thresholding yielded a quantitative viability assessment based on pixel-wise colour comparison. This provided single-cell resolution, as opposed to spectrophotometric assays. The same viability assay was also applied successfully to 3D culture yielding sharp cellular images from the surface (Supp. Fig. 8a), with the caveat that information from deeper than the superficial surface could not be obtained.

**Cell culture preparation** Human breast adenocarcinoma epithelial MCF-7 cell line was purchased from the American Type Culture Collection (ATCC). Cells were cultured in RPMI Medium 1640 (22400089, Gibco) supplemented with 10% fetal bovine serum (FBS-HI-12A, Capricorn Scientific), 100 $U/mL$ penicillin and 100 $\mu g/mL$ streptomycin (15140-122, Gibco) in 5% CO2 at $37°C$. Formation of spheroids was induced by seeding MCF-7 cells at 2x$10^4$ cells/well, in 96-well U-bottomed ultra-low attachment plates (174929, Life Technologies) with shaking for 2-3 days. 6-well plates (3516, Corning) were modified by drilling a 7 $mm$ circular window through the bottom of each well and having the quartz coverslip glued using liquid epoxy to cover the window. Cells were seeded directly on the modified 6-well plates and imaged directly on the quartz coverslip window.

### 2 Microtopography

#### 2.1 Isotropic response of fluorescent samples

FUSE is well suited to exploit the photometric stereo method, first introduced by Robert Woodham in 1980 [1], to obtain 3D surface microtopographical features (Supp. Fig. 1). Woodham's original assumptions require a Lambertian surface that perfectly diffused reflection. This is unrealistic for most real-life materials and use cases, which are non-Lambertian and anisotropic, which has led to multiple methods being developed to relax this requirement [2, 3]. The isotropic response [4] of fluorescently stained biological samples, in this case, is ideal. Since fluorescence emissions radiate isotropically with ideal Lambertian characteristics [5, 6] and the glass component of the objective is not DUV transmissive, errors associated with reflected DUV light are avoided, capturing a better and more accurate estimation of the 3D microtopography.

#### 2.2 Photometric Stereo

The photometric stereo method describes [7] the relationship between the luminous intensity ($I$) (or apparent brightness) to the surface irradiance ($I_0$), the cosine of the angle between the direction of the illumination source ($\hat{s}$) and the surface normal ($\hat{n}$), and the albedo (or reflection coefficient) as a constant factor ($A$).

$$I = A.I_0.\hat{s}.\hat{n} \tag{1}$$

With no specular reflections and cast shadows from the Lambertian fluorescent sample, each pixel is represented as,

$$m_i = \hat{s}_i.\vec{n} \tag{2}$$

where ($i$) is the light source coming from the vector direction ($\hat{s}_i$), with an intensity of ($m_i$), where $\vec{n} = [n_x, n_y, n_z]^T$ is a unitless vector of magnitude ($A.I_0$) and direction ($\hat{n}$). A sequence of 3 measurements can thus be expressed as,

$$\begin{bmatrix} S_x1 & S_y1 & S_z1 \\ S_x2 & S_y2 & S_z2 \\ S_x3 & S_y3 & S_z3 \end{bmatrix} \begin{bmatrix} n_x \\ n_y \\ n_z \end{bmatrix} = \begin{bmatrix} m_1 \\ m_2 \\ m_3 \end{bmatrix} \tag{3}$$

If the sample and the illumination sources are not on the same plane, and the illumination source matrix is non-singular, the system of linear equations can be solved for $\vec{n}$. For cases where there are more than 3 measurements, an estimation of the normal vector $\vec{n}$ can be made by minimising the residual error with the source vector, $\arg\min_{n_x,n_y,n_z} \sqrt{\sum_i (\hat{s}_i.\vec{n} - m_i)^2}$ Essentially, the overdetermined system of linear equations represents every pixel's intensity being proportionate to the dot product of the illumination vector and the surface normal. Thus, this allows the surface normals of every pixel of the sample to be obtained. The surface normals can be further interpreted with geometrical arguments to produce an approximate depth map.

### 3 Model development and analysis with CelebA

**Data** CelebA is a curated and processed dataset that contains the faces of celebrities. It is commonly used to benchmark deep learning models for single-image super-resolution. We used this dataset during the development of our model as a simple and sterile way of observing iterative improvements and can be compared with other state-of-the-art models. At the size of 218x178 *px*, the colour images in this dataset sit in a suitable zone where they are small enough to fit into memory (unlike our 4000x3000 *px* micrographs), but large enough to contain interesting and learnable features, and push the memory limits of large GPUs (unlike other datasets like CIFAR). We used 90% of the 202,599 faces for training for the model and the remaining 10% for comparative evaluation of the outputs.

**Performance measures** Four performance measures were used to provide a quantitative evaluation of the model (Supp. Fig. 9). The Learned Perceptual Image Patch Similarity (LPIPS) is a popular measure that aims to compare "perceptual" features that are robust to pixel-level photometric augmentations. The Structural Similarity Index (SSIM) compares images using hand-crafted aggregations of pixel-level information that empirically and intuitively reflect "luminance", "contrast", and "structure". Diversity is an empirical estimation of variation in generated images. It is computed as the sample-averaged pixel-wise standard deviation across samples. The final measure is simply the converged loss during model training, commonly used in previous studies with normalising flows. When considering the training objective of normalising flows is the minimisation of estimated likelihood, the loss is a direct measure of how well the learned distribution fits the data.

**Ablation study** FUSE-Flow is highly modular and studying each part independently enabled insights into the effects of the module on the overall model (Supp. Fig. 3). The image-encoding adaptive U-Net can be trained independently (with binary cross entropy loss) to serve as a baseline. It does a decent job of resolving well-defined facial features from the low-resolution versions. However, it displays some issues that are very characteristic of U-Nets trained with pixel-wise loss functions: very "average" and generic facial features, smoothed and undetailed surfaces (especially hair), and blurry edges. It also generates constant outputs since U-Nets are deterministic. When integrating the frozen pre-trained Adaptive U-Net to a normalising flow that is further trained with negative log-likelihood (NLL), not only are the generations now variable, but many of the downsides of U-Nets are now resolved. Edges are sharper, the texture is more detailed, and facial features are more personalised. The introduction of a self-attention mechanism like the Squeeze-and-Excitation module further improved the performance of the overall model. LPIPS improved significantly for FUSE-Flow compared to the original U-Net baseline, reflecting the observed visual improvement. SSIM, however, dropped. This is not entirely unexpected since U-Nets produce the most "average" result given the aleatoric uncertainty, which is the closest it can get to the ground truth without extra information not provided in the inputs. Conversely, each generated sample from FUSE-Flow includes some bias based on the sampled prior distribution that allows it to be high-resolution even though it lacks necessary information from the inputs. Diversity is drastically different unexpectedly, and the loss dropped compared to not using a self-attention mechanism.

**Controllable variation** There are two ways to control the generation of a trained FUSE-Flow. The first is through the input of a low-resolution image that conditions the generation. The other is by controlling the generative prior. It can be controlled either by changing the parameters of the prior (mean and standard deviation) or the way it is sampled. The standard deviation of the prior is often called "Temperature" or $\tau$ in prior studies using normalising flows and is a great way to manipulate the creativity of the model (Supp. Fig. 10a). Diversity is directly related to the Temperature, and we empirically demonstrated a clear and smooth relationship between them (Supp. Fig. 10b). As an aside, Supp. Fig. 10a demonstrates why SSIM decreases even though the detail of generated outputs improves. The model happened to assume curly hair was a possible ground truth, which unfortunately is very different to the actual ground truth (the reference).

### 4 Supplementary Bibliography

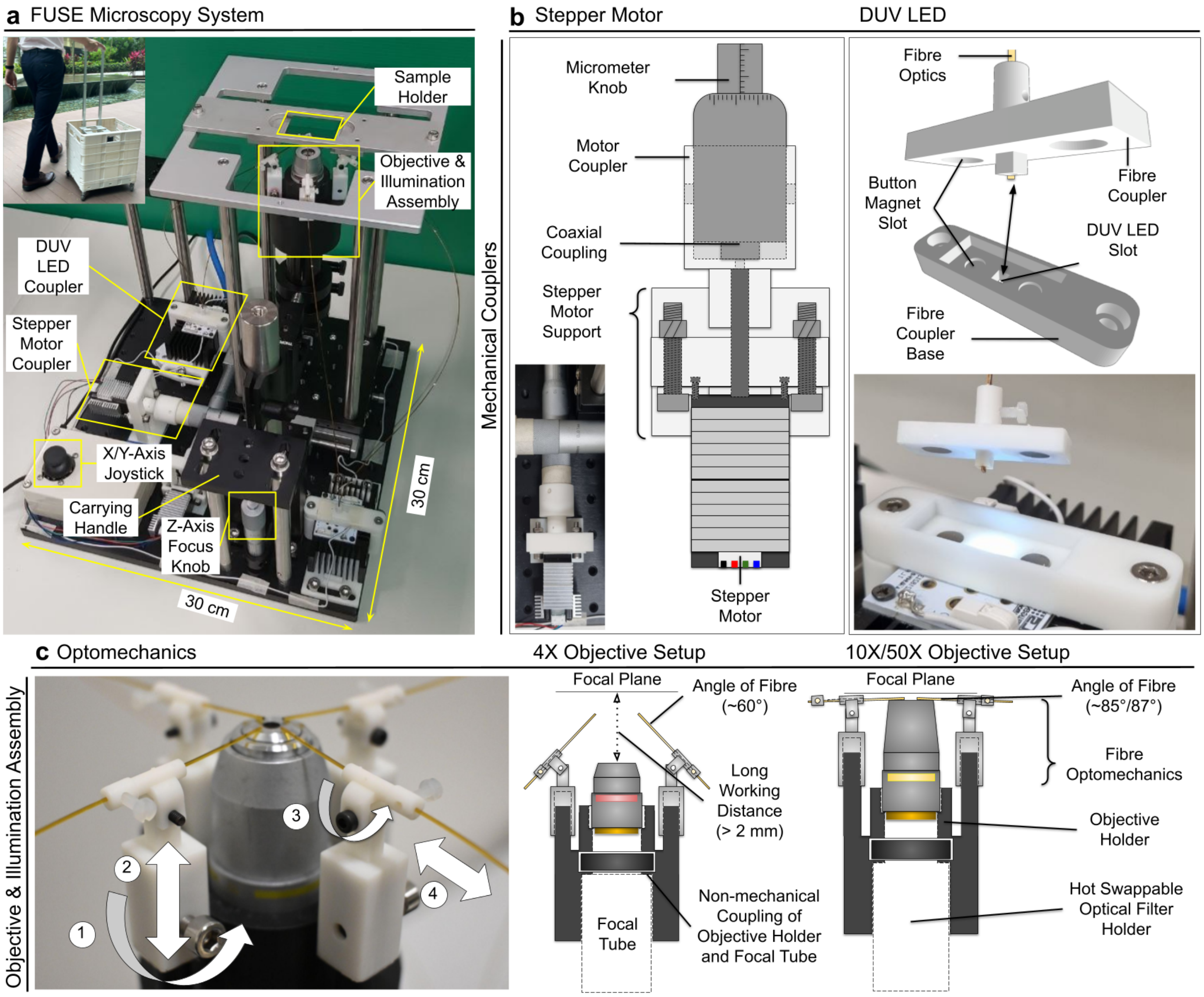


**Supp. Fig. 1: FUSE component details. a.** Compact and easily portable microscope with 30x30 *cm* footprint . **b.** Mechanical co-axial coupling of stepper motors to manual stage allowed for automation of FUSE. Fibre optics were magnetically butt coupled to DUV LEDs enabling DUV illumination to be transmitted to sample. **c.** Objective and illumination assembly comprised fibre optomechanics and objectives (4X/10X/50X). Fibre optomechanics allowed four degrees of freedom for optical fibres, enabling even illumination of samples and precise control over the direction and angle of illumination. Objective holders were non-mechanically coupled to focal tube, allowing quick swaps to be made to magnification.

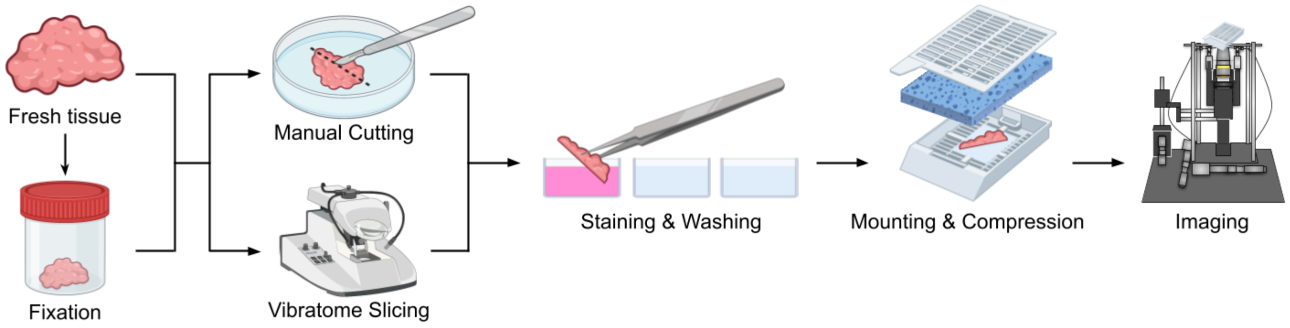


**Supp. Fig. 2: General tissue sample preparation.** Tissue was imaged either fresh or fixed by formalin. Tissue was manually trimmed with a blade or by vibratome slicing to expose *enface* surfaces for imaging. Desired pieces of tissue sample were stained in fluorescent dye concoction for 30-60 seconds. After staining, samples were washed twice in 1X PBS to remove excess stain. Rinsed samples were mounted onto modified histology cassettes and compressed with biopsy pads. After imaging, samples may be sent for downstream H&E processing for correspondence to FUSE images.

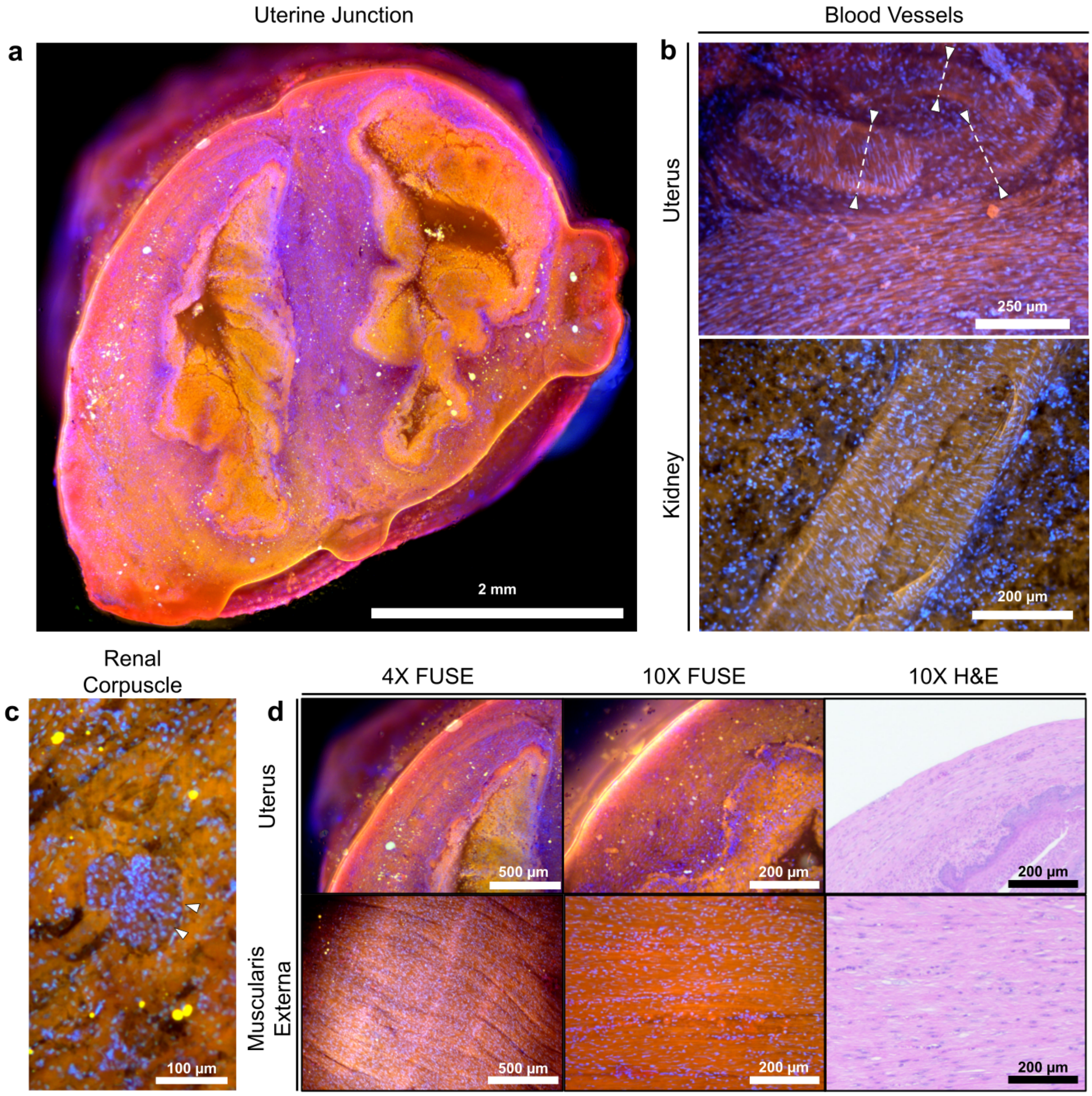


**Supp. Fig. 3: Histological features of murine organs. a.** Mosaiced image of cut fresh rat cervix revealed the merging of two uterine horns distal to uterine junction. Inner surface of endometrium was visibly textured (blue stain, nucleus stain Hoechst 33342; red-orange stain, counterstain rhodamine B). **b.** Above, three interlacing blood vessels were located in stratum vasculare of fresh rat uterus (10X). Vessels had approximate diameter of 140 µm, 130 µm, and 75 µm with a wall thickness of 25 µm, 25 µm, and 18 µm (left to right). Partially cut surface of a large blood vessel from fresh mouse kidney revealed inner surface of blood vessel (10X). **c.** Renal glomerulus in fresh mouse kidney shows epithelial parietal cells (white arrowheads) of Bowman's capsule (10X). **d.** 4X, 10X images, and corresponding H&E histology of fresh rat cervix showing lumen of a single uterine horn and muscularis externa. Nuclei of smooth muscle cells from outer longitudinal layer were clearly visualised at both 4X and 10X.

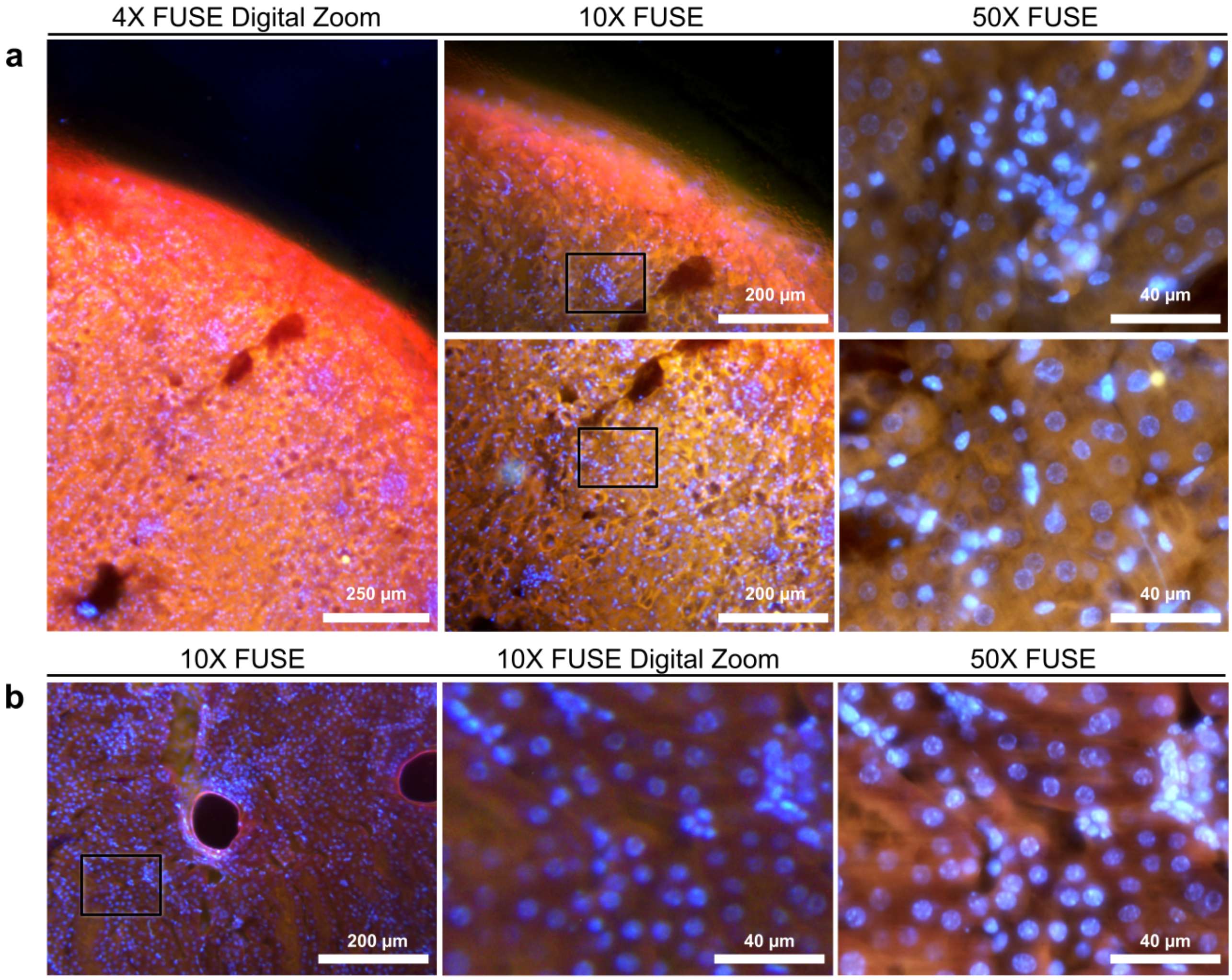


**Supp. Fig. 4: Thick tissue imaging at high magnification. a.** FUSE obtained multiscale (4X/10X/50X) images from thick (1 *mm*) fresh kidney samples. Rapid imaging of thick and unfixed samples could be conducted with lower magnifications (4X/10X) to quickly identify specific regions of interest before subsequent high-resolution imaging (50X) of important features. Distinct cell types were visible in renal corpuscle and renal tubules. **b.** High magnification (50X) imaging of thick fixed mouse kidneys illustrated increase in level of detail and resolution compared with lower magnification (10X). Sub-nuclear details could be identified from thick tissue samples with 50X FUSE.

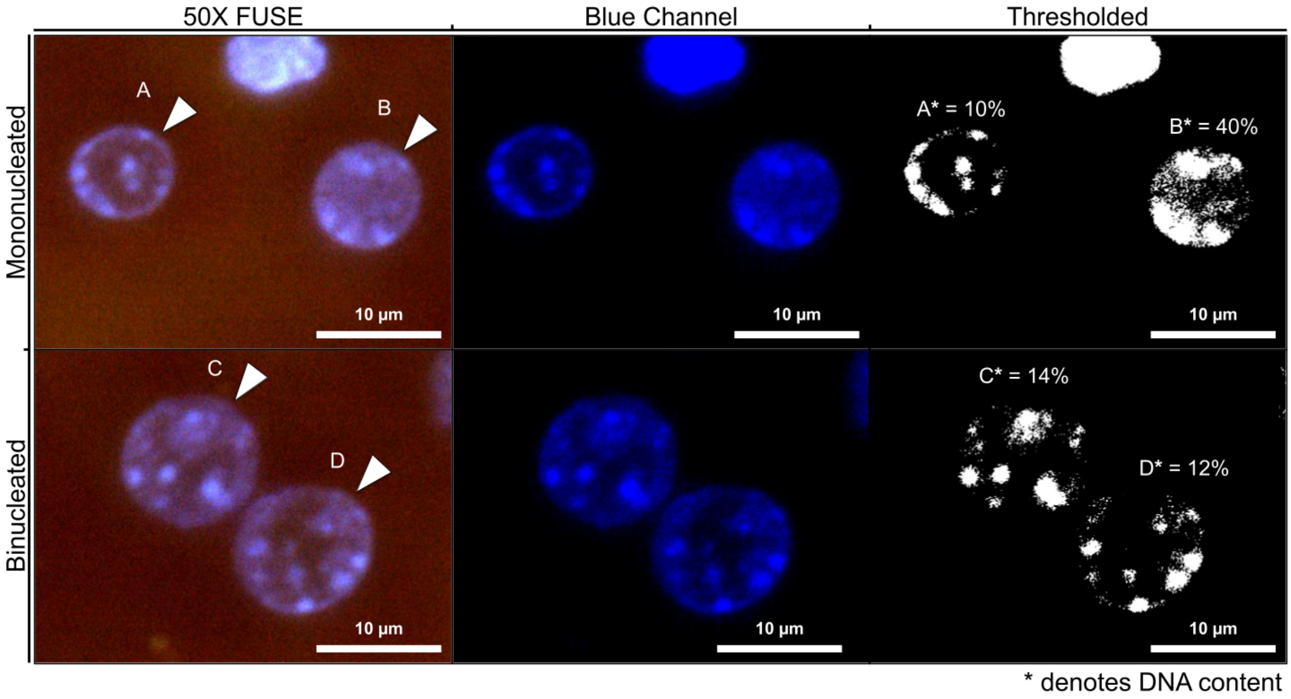


**Supp. Fig. 5: Cell cycle phase analysis through quantification of DNA content in thick samples.** Hepatocyte cell cycle phases were inferred by quantitative analysis of DNA content in hepatocyte nuclei from thick liver samples ($> 1\ mm$). 2 mononucleated hepatocyte nuclei (A & B) had significantly different DNA area expression within their nuclei envelope, suggesting significantly less DNA content in A than in B. A was inferred to be in the earlier Gap 1 (G1) phase, while B was in the later DNA synthesis (S) or Gap 2 (G2) phase. Similar nuclear sizes of A and B show that B was not in mitosis (M) phase, and the two nuclei were of the same ploidy class (2n). Binucleated hepatocyte nuclei (C & D) expressed similar DNA content, indicating similar cell cycle phases (G1 or early S), which was expected of binucleated hepatocytes.

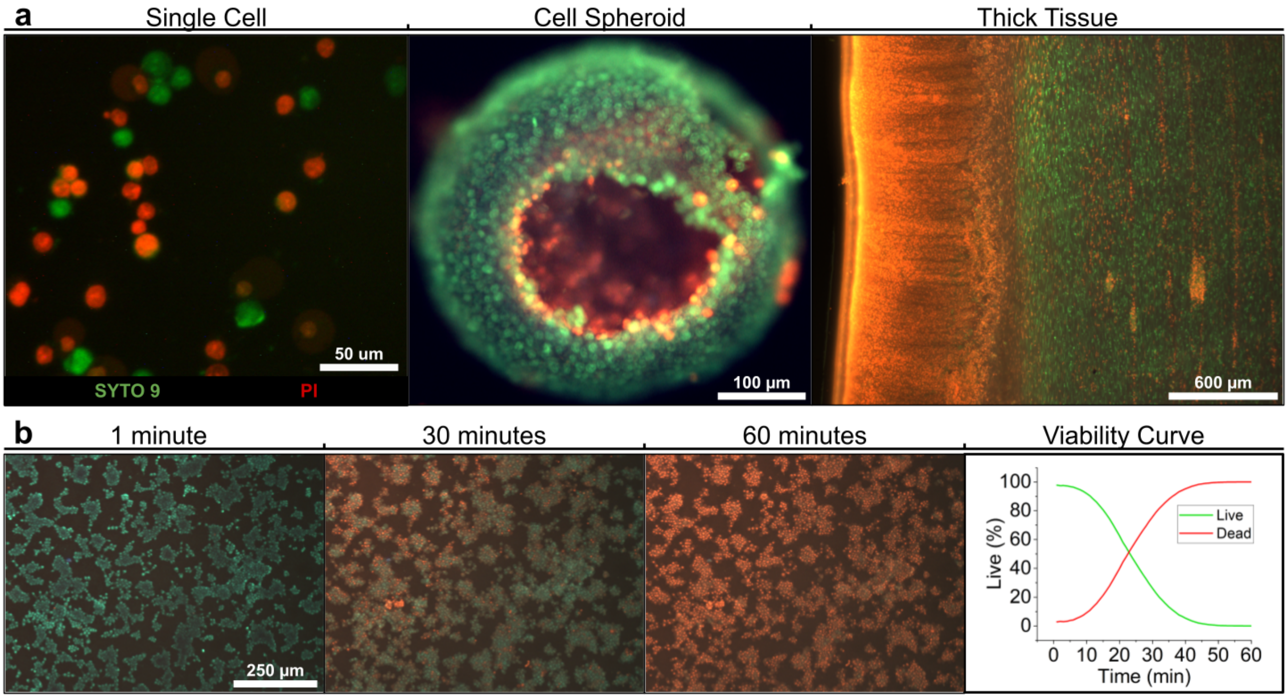


**Supp. Fig. 6: Viability from single cell to thick tissue. a.** Fluorescence cell viability assay (SYTO 9, propidium iodide) was effectively imaged with FUSE for 2D and 3D cell cultures and thick tissue samples. Spheroid (MCF-7) imaging showed a red necrotic core contrasting with the green proliferative layer. Viability staining highlighted tissue structure despite primarily staining cell nuclei. The mucosa, submucosa and muscularis mucosa of a fresh mouse stomach could be identified. **b.** Quantification of single cell (MCF-7) % viability over time after exposure to chemical insult (70% ethanol). Images were acquired at multiple time points over an hour. Cell segmentation and pixel-wise colour comparison produced quantitative viability assessments. Viability imaging on FUSE had single-cell spatial resolution, as opposed to intensity readouts from spectrophotometric assays.

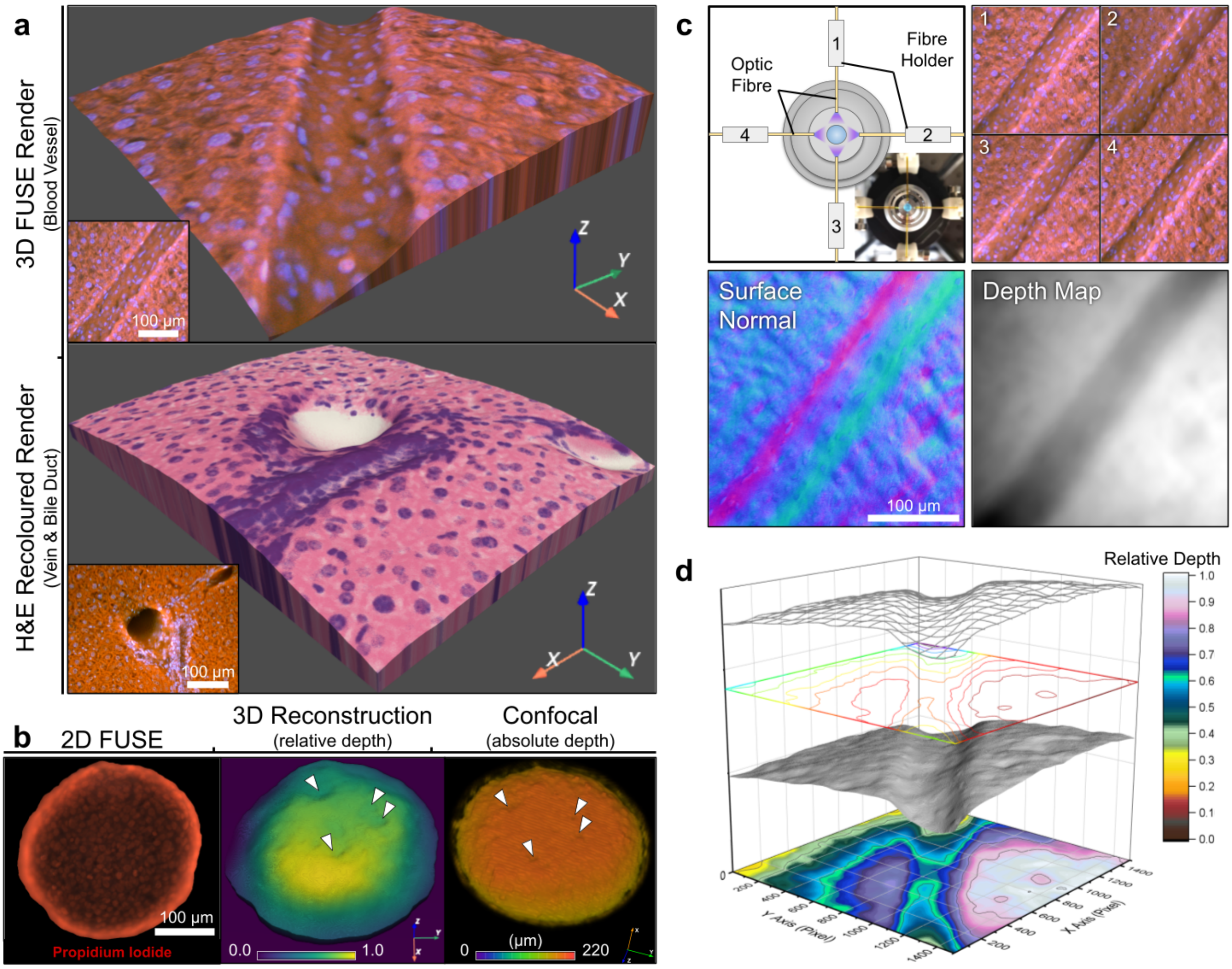


**Supp. Fig. 7: Fluorescence microtopography. a.** 3D visualisations of fixed mouse liver revealed intricate tubular structures, highlighting subtle surface texture details. Either original FUSE image or a virtual H&E image (among other variants) could be applied to colourise 3D model. **b.** 3D rendering of a cell spheroid, generated by photometric stereo, exhibited surface structures consistent with confocal imaging. While the algorithm intrinsically provided relative depth information, confocal microscopy produced absolute measurements. **c.** 3D renderings required the acquisition of four images, each illuminated by single fibres. The algorithm yielded the surface normal vector map, and subsequently generated depth map showed blood vessel's orientation, contours, and form. **d.** Variants of 3D topographical maps could be generated based on the depth map to suit specific needs.

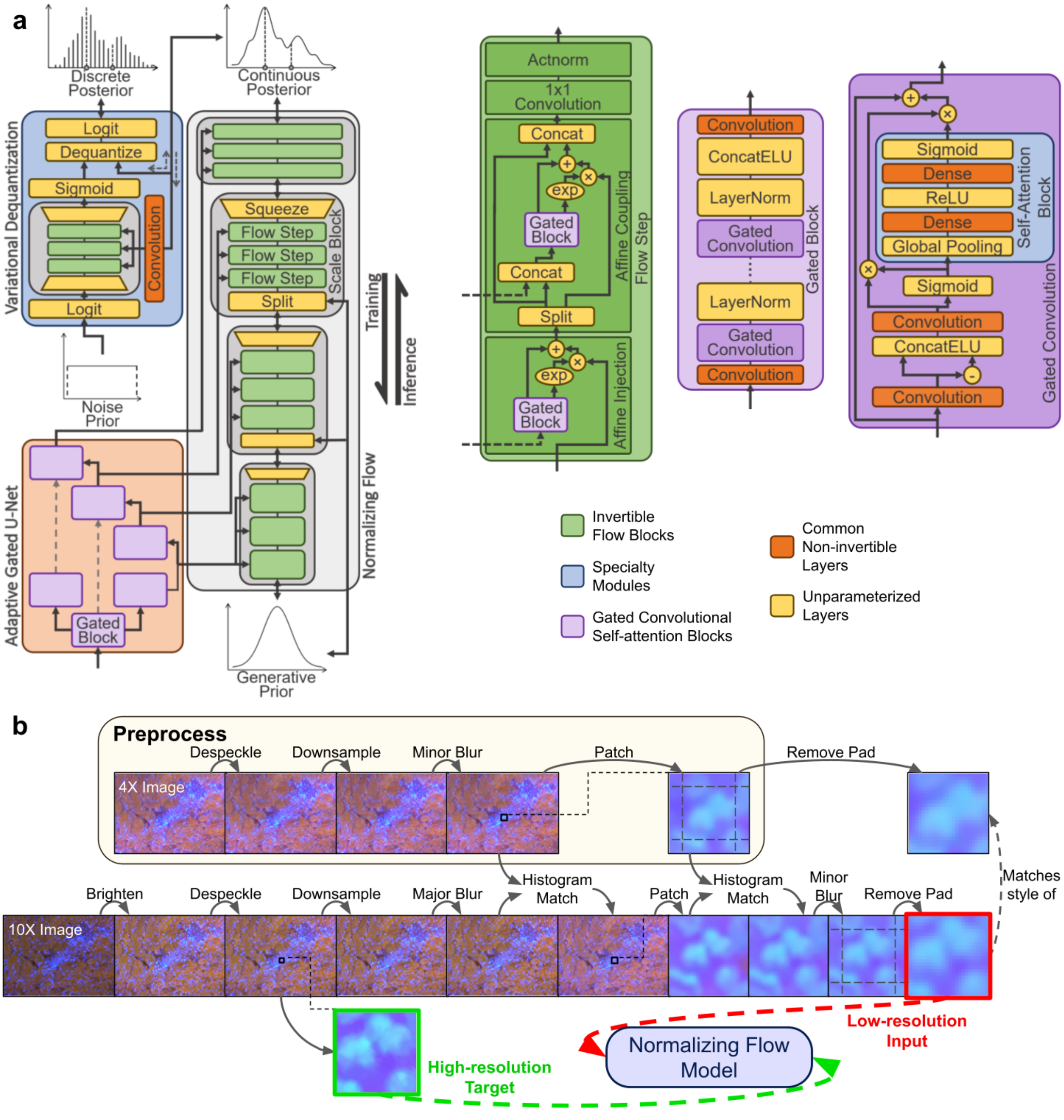


**Supp. Fig. 8: Detailed model architecture and data preparation pipeline. a.** FUSE-Flow consisted of three primary modules: normalising flow mapping between complex and simple continuous distribution, adaptive gated U-Net extracting image features to condition flow, and variational dequantisation module bridging discrete and continuous distributions via flexible noise addition (or subtraction). **b.** Data preparation for semi-supervised training approach entailed processing loosely aligned low and high-magnification image pairs to generate patches. Patches maintained pixel-accurate structural alignment with high-magnification image, yet emulated the style and resolution of low-magnification counterpart. After model training, data processing pipeline (titled "Preprocess") was employed on subsequent low-magnification images intended for enhancement.

**a**

| | LPIPS↓ | SSIM↑ | Diversity (σ)↑ | Convergent Loss↓ | |
|---|---|---|---|---|---|
| U-Net | 0.19678 | 0.68224 | 0 | 0.49828 | Binary Cross Entropy |
| U-Net + CBAM | 0.20487 | 0.66406 | 0 | 0.49799 | |
| U-Net + SE | 0.19024 | 0.69938 | 0 | 0.49657 | |
| U-Net + Flow | 0.11025 | 0.64891 | 6.7082 | 2.2528 | Bits per Dimension (scaled NLL) |
| U-Net + Flow + CBAM | 0.11578 | 0.64241 | 6.5283 | 2.2538 | |
| U-Net + Flow + SE **(FUSE-Flow)** | 0.11111 | 0.65159 | 6.4377 | 2.2270 | |

**b**

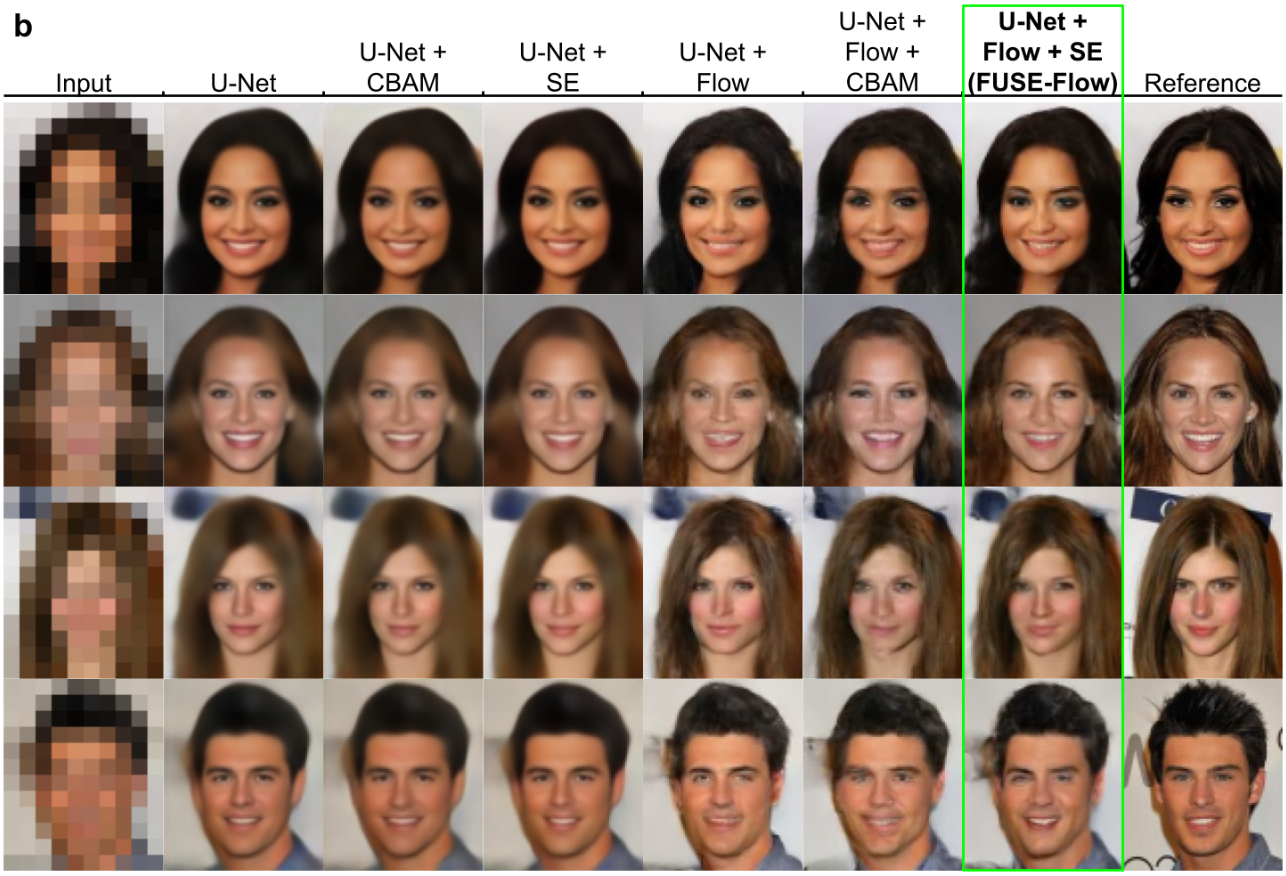


**Supp. Fig. 9: Ablation study on CelebA dataset. a.** Quantitative evaluation employed two common quality measures, Learned Perceptual Image Patch Similarity (LPIPS) and Structural Similarity (SSIM) metrics. Generative variation (diversity) was estimated using average pixel standard deviation ($\sigma$). Average loss was typically used for comparisons among models of the same class. **b.** The feature-extracting U-Net and the normalising flow are independently operable, with the U-Net being pre-trained and subsequently frozen during its use with the flow. Two self-attention mechanisms were tested and assessed—Convolutional Block Attention Module (CBAM) and Squeeze-and-Excitation (SE)—to improve the model's flexibility. Illustrative results from permutations of various modules are displayed.

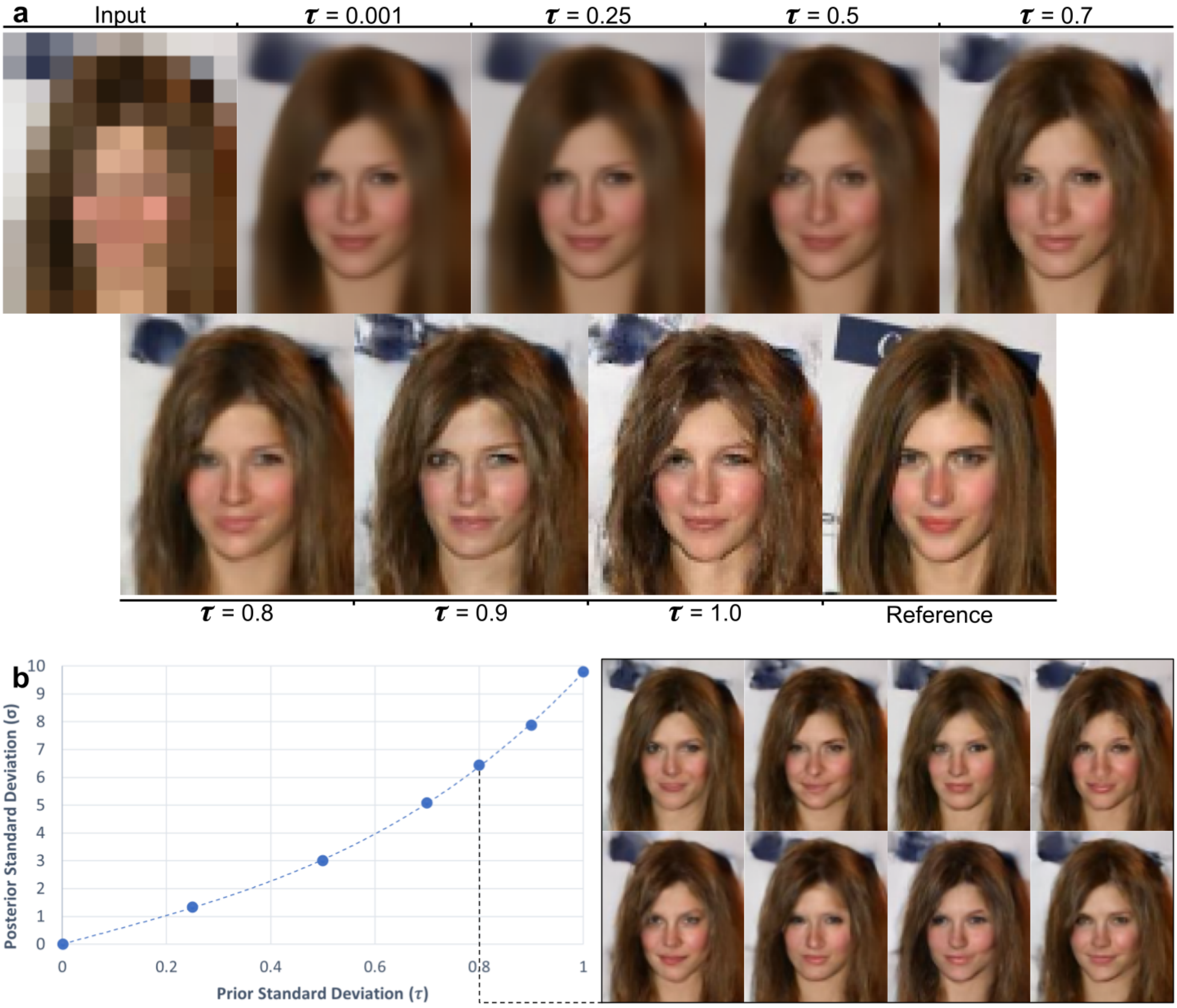


**Supp. Fig. 10: Overview of controllable variational capability. a.** Outputs at various temperature (user-defined standard deviation of Gaussian generative prior) levels highlight tradeoff between coarse resemblance to ground truth with deviating fine texture. **b.** There was a smooth relationship between the average pixel-wise standard deviation of the Gaussian generative prior ($\tau$) and generated images ($\sigma$). As $\tau$ increased, the diversity ($\sigma$) in generated faces increased.
